# Genetic dissection of the different roles of hypothalamic kisspeptin neurons in regulating female reproduction

**DOI:** 10.1101/464412

**Authors:** Luhong Wang, Charlotte Vanacker, Laura L. Burger, Tammy Barnes, Yatrik M. Shah, Martin G. Myers, Suzanne M. Moenter

## Abstract

The brain regulates fertility through gonadotropin-releasing hormone (GnRH) neurons. Estradiol induces negative feedback on pulsatile GnRH/luteinizing hormone (LH) release and positive feedback generating preovulatory GnRH/LH surges. Negative and positive feedback are postulated to be mediated by kisspeptin neurons in arcuate and anteroventral periventricular (AVPV) nuclei, respectively. Kisspeptin-specific ERα knockout mice exhibit disrupted LH pulses and surges. This knockout approach is neither location-specific nor temporally-controlled. We utilized CRISPR-Cas9 to disrupt ERα in adulthood. Mice with ERα disruption in AVPV kisspeptin neurons have typical reproductive cycles but blunted LH surges, associated with decreased excitability of these neurons. Mice with ERα knocked down in arcuate kisspeptin neurons showed disrupted cyclicity, associated with increased glutamatergic transmission to these neurons. These observations suggest activational effects of estradiol regulate surge generation and maintain cyclicity through AVPV and arcuate kisspeptin neurons, respectively, independent from its role in the development of hypothalamic kisspeptin neurons or puberty onset.

**Significant Statement:** The brain regulates fertility through gonadotropin-releasing hormone (GnRH) neurons. Ovarian estradiol regulates GnRH pulses (negative feedback) and the GnRH surge release that ultimately triggers ovulation (positive feedback). Kisspeptin neurons in the arcuate and anteroventral periventricular nuclei are postulated to convey negative and positive feedback to GnRH neurons, respectively. Kisspeptin-specific ERα knockout mice exhibited disrupted negative and positive feedback. However, it is not clear what roles each kisspeptin population plays, and not possible to separate their roles during development vs adulthood in this model. Here we utilized CRISPR-Cas9 to disrupt ERα in each population in adulthood. We found activational effects of estradiol regulate surge generation and maintain cyclicity through AVPV and arcuate kisspeptin neurons, respectively, independent from estradiol action during development.

## Introduction

Infertility is a common clinical problem affecting 15% of couples; ovulatory disorders account for 25% of this total (1). The hypothalamic-pituitary-gonadal axis controls reproduction and malfunction of this axis can cause ovulatory dysfunction and/or other disturbances of the reproductive cycle (2, 3). Gonadotropin-releasing hormone (GnRH) neurons form the final common pathway for central neural regulation of reproduction. GnRH stimulates the pituitary to secrete luteinizing hormone (LH) and follicle-stimulating hormone, which regulate gonadal steroid and gamete production. Estradiol, via estrogen receptor alpha (ERα), plays crucial roles in both homeostatic negative feedback and positive feedback action on GnRH/LH release in females (4–6). Low estradiol levels suppress pulsatile GnRH/LH release, whereas sustained elevations in estradiol during the late follicular phase of the cycle cause a switch of estradiol feedback action from negative to positive, inducing prolonged GnRH/LH surges, which ultimately triggers ovulation (7). As GnRH neurons typically do not express detectable ERα (8), estradiol feedback is likely transmitted to GnRH neurons by ERα-expressing afferents.

Kisspeptin neurons in the arcuate and anteroventral periventricular (AVPV) regions are estradiol-sensitive GnRH afferents that are postulated to mediate estradiol negative and positive feedback, respectively (9, 10). Kisspeptin potently stimulates GnRH neurons and *Kiss1* mRNA is differentially regulated in these nuclei by estradiol (10–16). ERa in kisspeptin cells is critical for estradiol negative and positive feedback, as kisspeptin-specific ERα knockout (KERKO) mice exhibit higher frequency LH pulses and fail to exhibit estradiol-induced LH surges(17–20).

Although informative, the KERKO model has several caveats that limit interpretation. First, ERα is deleted as soon as *Kiss1* is expressed, before birth in arcuate kisspeptin neurons (also called KNDy neurons for coexpression of kisspeptin, neurokinin B and dynorphin) and before puberty in AVPV kisspeptin neurons (21, 22). This may cause developmental changes in these cells and/or their networks. Second, ERα is deleted from all kisspeptin cells, thus making it impossible to assess independently the role of AVPV and arcuate kisspeptin neurons.

Combining CRISPR-Cas9 with targeted viral vector injection allows deletion of ERα in a nucleus-specific and temporally-controlled manner to address the above caveats (23). We designed cre-dependent AAV vectors that carry single guide RNAs (sgRNAs) that target *Esr1* (encoding ERα) or *lacZ* and delivered these vectors to the AVPV or arcuate of adult female mice that express Cas9 in kisspeptin cells. We then compared the reproductive phenotypes as well as kisspeptin neuronal physiology in AAV-*Esr1* vs AAV-*lacZ* targeted mice and KERKO mice.

## Results

### AVPV kisspeptin neurons from KERKO mice exhibit decreased firing rate and excitability

*compared to controls and fail to respond to estradiol*. We first used extracellular recordings to monitor the spontaneous firing rate of YFP-identified AVPV kisspeptin neurons in coronal brain slices from ovary-intact control and KERKO mice. As the persistent cornified vaginal cytology of KERKO mice is similar to that observed during estrus (19), we used mice in the estrous stage of the reproductive cycle as controls. The firing frequency of AVPV kisspeptin neurons was lower in ovary-intact KERKO mice compared to controls (Fig 1A, B, two-way ANOVA/Holm-Sidak, p=0.0001). To test if the firing rate of AVPV kisspeptin neurons in KERKO mice responds to circulating estradiol, we repeated this study in ovariectomized (OVX) mice and OVX mice with an estradiol implant producing constant physiologic levels (OVX+E)(24). OVX reduced and estradiol treatment increased firing rate in cells from control, but not KERKO, mice (Fig 1A, B two-way ANOVA/Holm-Sidak, control intact vs OVX p=0.009, intact vs OVX+E, p=0.02, OVX vs OVX+E, p<0.0001). As a result of this difference, the firing frequency is higher in cells from OVX+E control than OVX+E KERKO mice (p<0.0001). Statistical test parameters for all figures are in SI Appendix Table S1.

**Fig 1.**
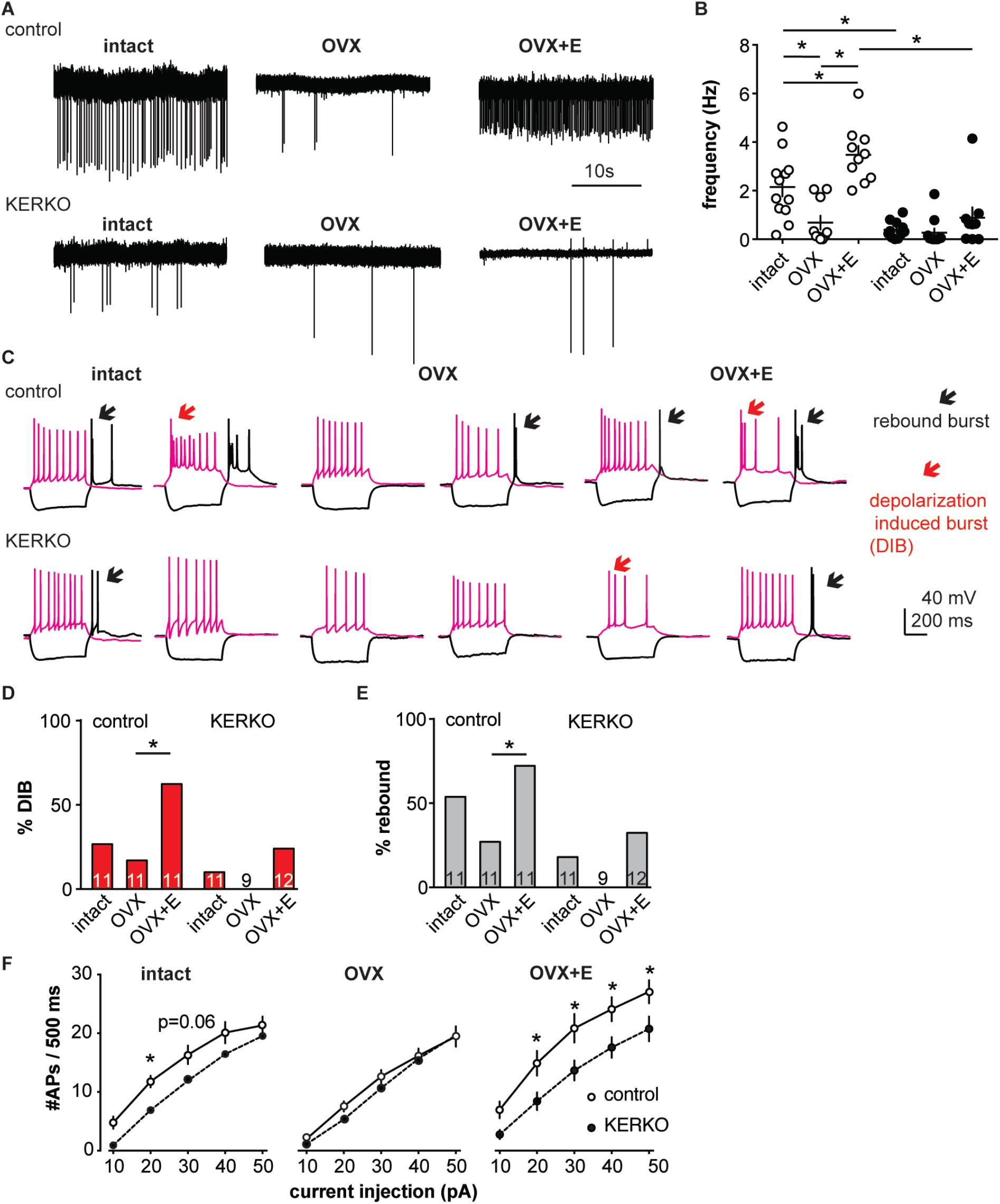
AVPV kisspeptin neurons from KERKO mice are less excitable compared to those from control mice and are not regulated by estradiol. (A), representative extracellular recordings for cells from control and KERKO mice from ovary-intact, OVX and OVX+E groups. (B), individual values and mean±SEM firing frequency of cells from control (white circles, intact n=12, OVX n=10, OVX+E, n=10) and KERKO groups (black circles, intact n=11, OVX n=11, OVX+E, n=9). (C), representative depolarizing (magenta, +20pA, 500ms) and hyperpolarizing (black, −20pA, 500ms) firing signatures for cells from control and KERKO mice in ovary-intact (left), OVX (middle) and OVX+E (right) groups; black arrows indicate rebound bursts and magenta arrows indicate depolarization-induced bursts (DIB). Initial membrane potential was 70±2mV. (D-E), percent of cells exhibiting DIB (D) or rebound (E) bursts; cells per group is shown within the bar. (F), input-output curves for cells from control and KERKO mice; ovary-intact (left), OVX (middle) and OVX+E (right). * p<0.05

We next recorded the whole-cell firing signatures of neurons in these six groups in response to current injection. AVPV kisspeptin neurons in control mice exhibit a greater number of depolarization-induced bursts (DIB) and rebound bursts when estradiol is elevated, confirming previous observations (25) (Fig 1 C, D, E, Chi-square, DIB, p=0.02; rebound, p=0.02; Fisher’s exact *post hoc* test, DIB, OVX vs OVX+E, p=0.008, rebound OVX vs OVX+E p=0.03, for other paired comparisons, p>0.2)). In KERKO mice, these two types of bursts were rare (<25% of cells) in all steroid conditions tested and were not regulated by estradiol (Fig 1C, D, E, Chi-square, DIB, p=0.4, rebound, p=0.3). We also compared the action potential output of these cells in response to current injection (0-50 pA, 10 pA increments, 500 ms). Cells from ovary-intact KERKO mice generated fewer action potentials compared to controls. Action potential generation as a function of current injection was similar in cells from OVX control and OVX KERKO mice but was increased by estradiol only in control mice (Fig 1F, two-way repeated-measures ANOVA/Holm-Sidak, intact, 20 pA, p=0.03, 30 pA, p=0.06; OVX+E, 20 pA to 50 pA, p≤0.04). Reduced action potential firing of AVPV kisspeptin neurons from KERKO mice may be attributable at least in part to decreased input resistance compared to controls (SI Appendix Fig S1A, two-way ANOVA/Holm-Sidak control vs KERKO, intact, p=0.006, OVX, p=0.7, OVX+E p=0.02).

As both depolarization-induced bursts and rebound bursts are sensitive to NiCl (100 µM) (26) at levels that fairly specifically block T-type calcium channels, we measured T-type (I_T_) current density and voltage dependence. I_T_ current density was decreased in AVPV kisspeptin cells from gonad-intact KERKO mice compared to controls (SI Appendix, Fig S1 A, B, two-way repeated-measures ANOVA/Holm-Sidak, −50 mV, p=0.003; −40 mV, p=0.002; −30 mV, p=0.003). The voltage dependence of activation was not different between groups, but the voltage dependence of inactivation was depolarized in cells from KERKO mice (SI Appendix Fig S1 C, control vs KERKO, two-tailed unpaired Student’s *t*-test, V_1/2_ activation −52.2±1.6 vs −48.6±1.4 mV, p>0.1; slope 5.5±0.6 vs 5.5±0.7, p>0.1; V_1/2_ inactivation −74.8±4.1 vs −61.9±3.1 mV, p=0.03; slope −3.1±0.5 vs −4.2±0.3, p=0.1).

### Design and validation of sgRNAs that target *Esr1*

A caveat of studying the role of ERα in AVPV kisspeptin neurons using KERKO mice is that the deletion of ERα (encoded by *Esr1*) using cre recombinase under the control of the kisspeptin promoter is neither time-nor location-specific. We utilized the CRIPSR-Cas9 approach to achieve temporal and spatial control of *Esr1* gene knockdown. Two sgRNAs were designed that target exon1 of *Esr1* based on software prediction (27) and the efficiency of each guide tested *in vitro* in C2C12 mouse myoblast cells (28). The sgRNAs that target *Esr1* and a sgRNA that targets *lacZ* as a control were subcloned into the lentiCRISPRv2 plasmid (29), from which Cas9 and the sgRNA are expressed after transfection of C2C12 cells. Puromycin was used to select construct-expressing cells. After a ~4-week selection period, DNA was harvested and the *Esr1* region sequenced. Cells expressing either of the sgRNAs targeting *Esr1*, but not *lacZ*, exhibited a peak-on-peak sequencing pattern, indicating disruption of the gene (Fig 2A). As these *in vitro* experiments suggested these sgRNAs were able to mutate *Esr1*, we designed Cre-dependent AAV vectors to express each sgRNA and mCherry (to indicate infected cells) under control of the U6 promoter (Fig 2B). The AAV vector was bilaterally stereotaxically injected into the AVPV region of adult female mice that express Cas9 and GFP under control of the kisspeptin promoter (Fig 2C, SI Appendix Fig S3A); these groups are referred to as AVPV-AAV-*Esr1* or AVPV-AAV-*lacZ*. Only one guide was injected per animal to allow comparison of phenotypes when different areas of *Esr1* were targeted. The ERα knockdown efficiency of the two sgRNAs target *Esr1* was comparable. The infection rate for AVPV-AAV-*Esr1* was 81±4% (Fig 2D, *Esr1*-guide 1 [g1] 82±2%, n=3; *Esr1*-guide 2 [g2] 81±8%, n=3,) and only 28±1% of AVPV kisspeptin cells expressed ERα post infection (Fig 2D, n=3, *Esr1*-guide1 27±0.4%, n=3 *Esr1*-guide2 28±2%). In mice that received AVPV-AAV-*lacZ* (n=3), the infection rate was comparable at 82±2%, but there was no disruption of ERα; 72±2% of AVPV kisspeptin neurons expressed ERα, which is similar to control mice (11).

**Fig 2.**
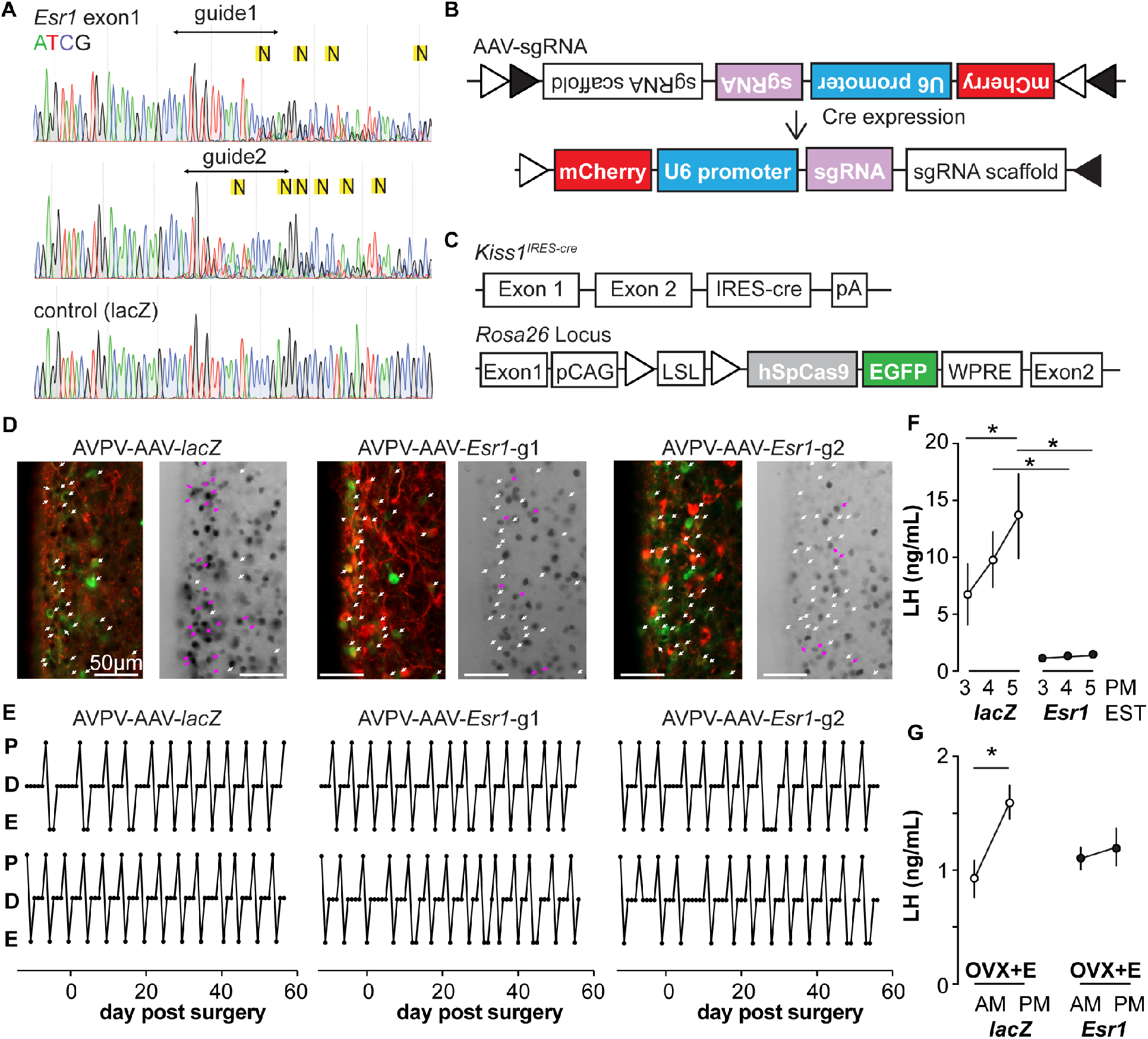
*In vitro* and *in vivo* validation of AVPV-AAV-*Esr1* guides. (A), sequencing from C2C12 cells transiently transfected with lentiCRISPR v2 with sgRNAs targeting *Esr1* (guide 1 [g1] top, guide 2 [g2] middle) or lacZ. N in yellow highlight indicates peak on peak mutations. (B-C), Schematic representation of (B) the Cre-inducible AAV vector delivering sgRNAs and (C) *Kiss1-*cre *Cas9-Gfp* mice. (D), AVPV-AAV-*lacZ*, -*Esr1* g1 or g2 were bilaterally delivered to the AVPV region (see SI Appendix Fig S3 A). Brain sections were processed to detect GFP (green), mCherry (red) and ERα (black), dual GFP/mCherry detection indicates infection of kisspeptin neuron (white arrows, left panel of each pair). AVPV-AAV-*Esr1* infected AVPV kisspeptin neurons exhibit decreased ERα expression compared to AVPV-AAV-*lacZ* infected cells (right panel of each pair, white arrows indicate ERα-negative, magenta arrows indicate ERα-positive infected cells). (E), representative reproductive cycles of mice that received AAV-*lacZ, g1 or g2*; E, estrus, D, diestrus, P proestrus, day 0 is the day of stereotaxic surgery. (F), mean±SEM proestrous LH surge measured at 3, 4, and 5 pm EST in AVPV-AAV-*lacZ* and AVPV-AAV-*Esr1* mice (mice receiving g1 or g2 combined). (G), mean±SEM estradiol-induced LH surge measured at 9 am and 5 pm EST from AAV-*lacZ* and AAV-*Esr1* OVX+E mice (mice receiving g1 or g2 were combined).

### Deletion of ERα in AVPV kisspeptin neurons in adulthood does not affect estrous cycles but disrupts preovulatory and estradiol-induced LH surges

We monitored the reproductive cycles of the mice injected with AAV-sgRNAs in the AVPV 12 days before and for up to eight weeks following surgery. Neither AVPV-AAV-*Esr1* guide (tested independently) nor the AVPV-AAV-*lacZ* disrupted reproductive cyclicity (Fig 2E), even in mice with a high rate of bilateral infection (~80%). These mice entered proestrus at the same frequency in the last 4 weeks as during the first 4 weeks (2 weeks pre-surgery + first 2 weeks post-surgery, two-way repeated-measures ANOVA/Holm-Sidak, before vs after, g1, n=3, 1.3±0.1 vs 1.6±0.1; g2, n=4, 1.2±0.1 1.4±0.1, *lacZ* n=4, 1.3±0.2 vs1.3±0.1, p>0.1 for each paired comparison). To test for the occurrence of estradiol positive feedback, we monitored both proestrous (preovulatory) and estradiol-induced LH surges in these mice. Surge data were similar for guide 1 and guide 2 and data from both guides were combined for group comparisons. Both proestrous and estradiol-induced LH surges were blunted after ERα knockdown (Fig 2F, G, two-way repeated-measures ANOVA/Holm-Sidak; G, *lacZ*, 3pm vs 5pm, p=0.04; *lacZ* vs *Esr1*, 4pm, p=0.006, 5 pm, p<0.0001, H, *lacZ* AM vs PM, p<0.0001).

### Decreased excitability of AVPV kisspeptin neurons in AAV-*Esr1* knockdown mice

To test if knockdown of ERα in adult AVPV kisspeptin neurons alters their intrinsic excitability, we recorded firing signatures of infected and uninfected cells in brain slices from AAV injected OVX+E mice. We again observed no difference between AVPV-AAV-*Esr1* g1 vs g2 and combined these data. Some cells were loaded with neurobiotin during recording for identification and ERa protein detected *post hoc* with immunofluorescence (Fig 3A, C, and IF *post hoc* portions of 3E-J). Cells not infected with AVPV-AAV-*Esr1* and cells infected with either AVPV-AAV-*Esr1* guide but in which ERα protein was detected exhibited similar firing signatures in terms of DIB and rebound bursts (Fig 3B, E, F). In contrast, cells infected by AVPV-AAV-*Esr1* that had undetectable ERα protein had reduced burst firing compared to AVPV-AAV-*lacZ* or uninfected groups (Fig 3E, F, Chi-square, DIB, p=0.008, rebound bursts, p=0.0008; Fisher’s exact *post hoc* test, DIB, *Esr1* vs *lacZ*, or vs uninfected, p≤0.03; rebound, *Esr1* vs *lacZ*, or vs uninfected p≤0.04; for other paired comparisons, p>0.5). The firing signature of AAV-*Esr1* infected cells with successful deletion of ERα was comparable to cells from KERKO mice (Chi-square, p>0.9 for both DIB and rebound bursts). Cells that lost detectable ERα after AVPVAAV-*Esr1* infection also produced fewer action potentials with current injection than cells infected with AAV-*lacZ* (Fig 3D left and center, two-way repeated-measures ANOVA/Holm-Sidak, *Esr1* vs *lacZ*, 20 pA, p=0.08; 30 to 50pA p≤0.02). This difference is not attributable to passive properties (SI Appendix Fig S2 A, B). The relationship between current injection and number of action potentials fired (input-output curve) in cells from KERKO and in AVPV-AAV-*Esr1* knockdown mice was only different at the highest level of current injected, with AVPVAAV-*Esr1* infected cells being less excitable (Fig 3D, right, two-way repeated-measures ANOVA/Holm-Sidak, 50pA, p=0.01), despite no change in input resistance (SI Appendix Fig S2 A,B KERKO vs AAV, p=0.08). Action potential properties from AVPV-AAV-*Esr1* knockdown cells with AVPV-AAV-*lacZ* control also differed. Specifically, loss of ERα led to decreased action potential rate of rise, a trend towards prolonged full-width half-maximum (FWHM), and hyperpolarized afterhyperpolarization potential (AHP) (Fig 3G, H, J, two-tailed unpaired Student’s *t*-test, H, p=0.02, J, p=0.0002; I, Mann-Whitney U-test, p=0.06).

**Fig 3.**
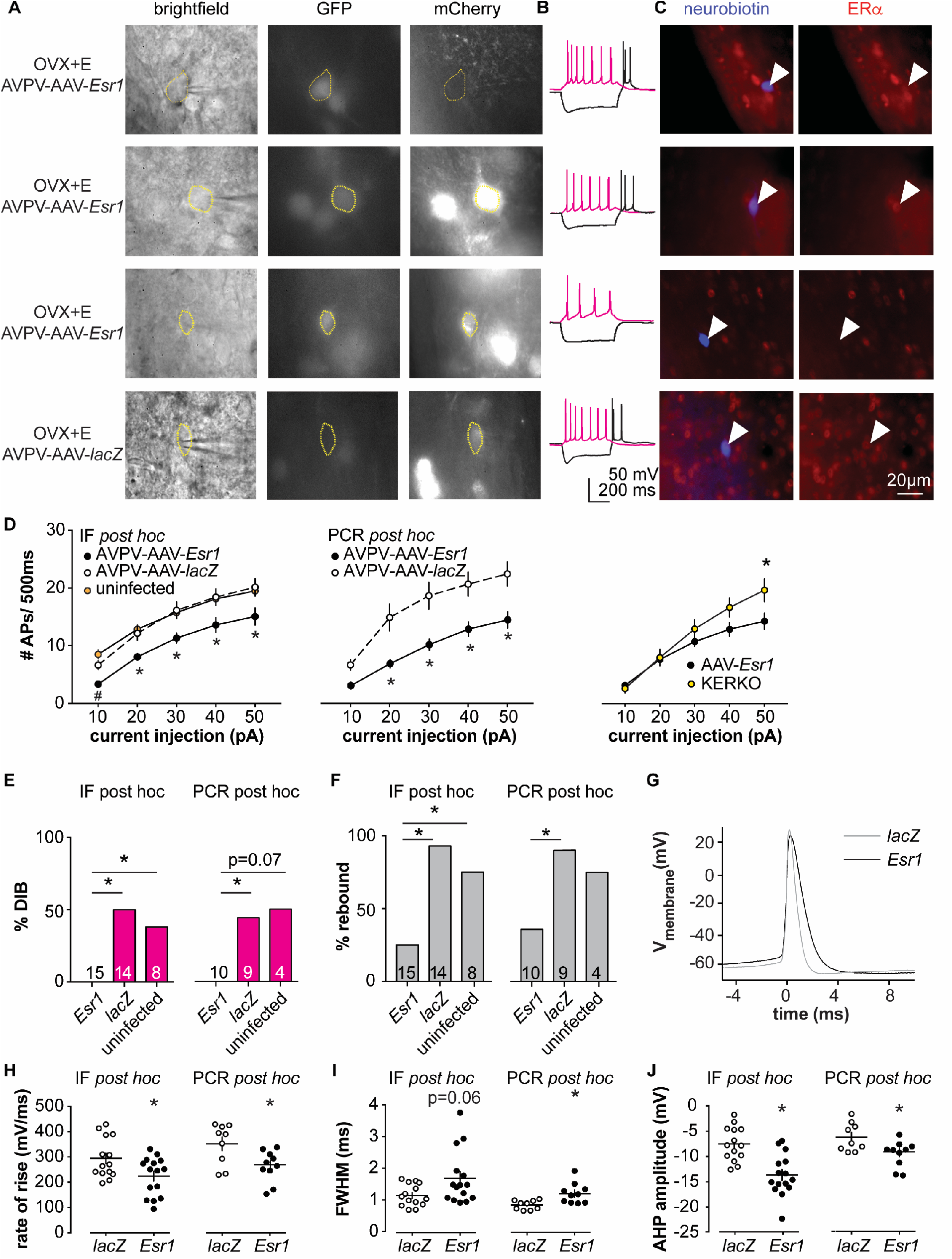
Decreased excitability of AVPV kisspeptin neurons in AVPV-AAV-*Esr1* knockdown mice. (A-C), whole-cell recording and immunofluorescence (IF) *post hoc* identification of ERα in recorded cells in OVX+E AVPV-AAV-*Esr1* infected mice. (A), visualization during recording; (B), representative depolarizing (+20pA, magenta) and hyperpolarizing (−20pA, black) firing signatures. (C), neurobiotin (blue) and ERα (red) staining after photobleaching of GFP and mCherry signals. From top to bottom: cells not infected by AVPV-AAV-*Esr1* and immunopositive for ERα; cells infected by AVPV-AAV-*Esr1* but still immunopositive for ERα; cells infected by AAV-*Esr1* and not immunopositive for ERα; cells infected by AVPV-AAV-*LacZ* and immunopositive for ERα. (D), left, input-output curves of infected cells with undetectable ERα in AAV-*Esr1* (third row in a-c, black circle, n=15), cells infected by *AVPV-*AAV-*lacZ* (bottom row in a-c, orange circle, n=14), and cells not infected by AAV (top row in a-c, white circle, n=8); middle, input-output curves from a separate set of cells in which *Esr1* status was confirmed by single-cell qPCR *post hoc* (AAV-*Esr1* black circle, n=10, AAV-l*acZ*, white circle, n=9); right, input-output curve of AVPV-AAV-*Esr1* knockdown (black circle) vs KERKO (yellow circle) cells. (E-F), percent of cells exhibiting DIB (E) or rebound bursts (F). Cells per group is shown within or on top of the bar. g, representative action potentials at the rheobase from *lacZ* vs *Esr1* infected cells. (H-J), individual values and mean±SEM rate of rise (H), full width at half maximum (FWHM, i) and afterhyperpolarization potential amplitude (AHP, j). * p<0.05 vs all other groups; # p<-.05 vs uninfected.

In parallel, we performed whole-cell patch-clamp recording with single-cell PCR *post hoc* identification of *Esr1* mRNA on a separate set of cells (AVPV-AAV-*Esr1* 10 cells from 4 mice, AVPV-AAV-*lacZ*, 9 cells from 3 mice). A similar decrease in burst firing and action potential input-output curve was observed in *Esr1* mRNA negative cells as was observed in cells verified to have undetectable ERα protein by immunofluorescence (Fig 3E, F, Chi-square, DIB, p=0.04, rebound p=0.03; Fisher’s exact *post hoc* test, DIB, *Esr1* vs *lacZ* p=0.03. *Esr1* vs uninfected, p=0.07; rebound, *Esr1* vs *lacZ* p=0.02; for other paired comparisons, p>0.2). Absence of *Esr1* mRNA expression was again associated with decreased number of action potentials in response to current injection (Fig 3E middle, *Esr1* vs *lacZ*, p<0.002 for 20 to 50pA steps). Absence of *Esr1* mRNA, similar to loss of ERα protein, led to decreased action potential rate of rise, prolonged FWHM, and AHP (Fig 3G-J right, two-tailed unpaired Student’s *t*-test, H, p=0.02; I, p=0.006, J, p=0.02). Single-cell PCR analysis also indicates that a lower percent of AVPV-AAV-*Esr1* knockdown cells express *Kiss1* and a trend to increase in *Esr2* mRNA (AVPV-AAV-*Esr1* 23 cells from 4 mice, AVPV-AAV-*lacZ*, 16 cells from 3 mice, SI Appendix Fig S4). Interestingly, expression of the mRNA for progesterone receptor (*Pgr)* did not differ between groups (SI Appendix Fig S4), suggesting the estradiol-dependence of this gene may be paracrine regulated in the brain as in other tissues (30). We also examined gene expression for several ion channels, but none showed any changes or patterns of expression among groups (SI Appendix Fig S4).

### Deletion of ERα in arcuate kisspeptin neurons in adulthood disrupted estrous cycles

To examine the role of estradiol feedback on arcuate kisspeptin neurons, we delivered the AAV-sgRNAs bilaterally to the arcuate region to knockdown ERα in these cells (SI Appendix Fig S3 B); these groups are referred to as Arc-AAV-*Esr1* or Arc-AAV-*lacZ*. The infection rate for Arc-AAV-*Esr1* was 92±3% (SI Appendix Fig S3B, n=3, *Esr1*-guide1 96±2%, n=3 *Esr1*-guide2 86±2%) and only 34±3% of KNDy neurons expressed ERα post infection (Fig 4A, n=3, *Esr1*-g1 38±0.3%, n=3 *Esr1*-g2 30±5%). In mice that received Arc-AAV-*lacZ*, the infection rate was comparable (Fig 4A, n=3 94±3%), and 92±1% of KNDy neurons expressed ERα, similar to control mice(11). Reproductive cycles were monitored for 12 days before and for up to eight weeks following surgery. In contrast to mice with Arc-AAV-*Esr1* targeted to the AVPV region, mice with the same virus targeted to the arcuate began exhibiting disrupted cyclicity three to four weeks post surgery (Fig 4B). These mice entered proestrus less frequently after surgery than before (two weeks pre-surgery + first two weeks post-surgery, Fig 4D, two-way repeated-measures ANOVA/Holm-Sidak, g1, p=0.002, g2, p=0.03). There was no difference in LH pulse frequency measured on the day of estrus or mean levels between Arc-AAV-*Esr1* vs Arc-AAV-*lacZ* injected mice on estrus (Fig 4C, E, F). Notably, LH response to kisspeptin and to GnRH was reduced in Arc-AAV-*Esr1* mice (Fig 4G, two-way repeated measures ANOVA/Holm-Sidak, *lacZ*, control vs kisspeptin or GnRH, p<0.001; *Esr1* vs *lacZ* for kisspeptin and GnRH both, p≤0.002).

**Fig 4.**
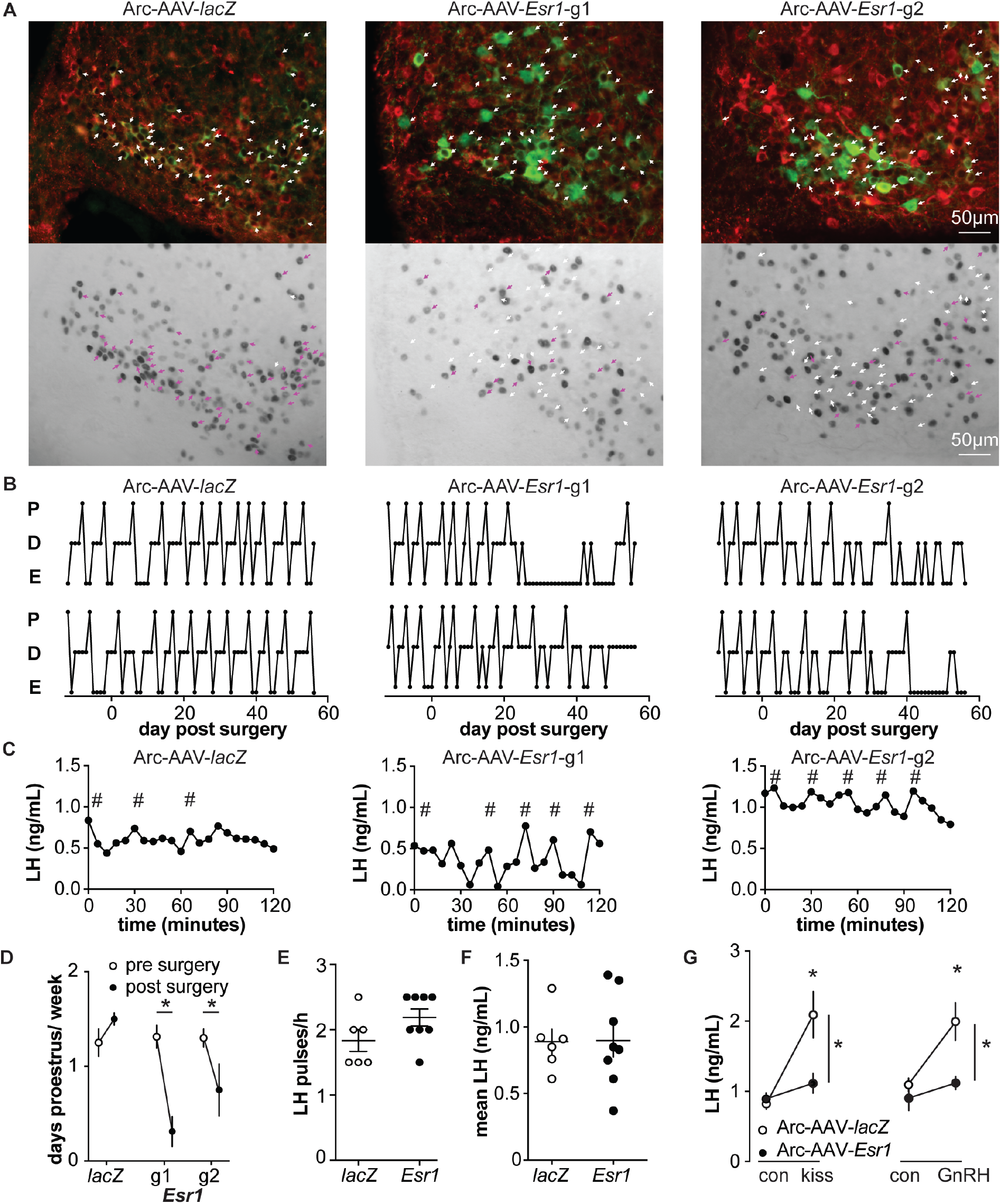
Deletion of ER in arcuate kisspeptin neurons (A), Arc-AAV-*lacZ* and Arc-AAV-*Esr1* (g1 or g2) were bilaterally delivered to arcuate region (see SI Appendix Fig S3 B). Brain sections were processed to detect GFP (green), mCherry (red) and ERα (black). Arc-AAV-*Esr1* infected arcuate kisspeptin neurons exhibit decreased ERα expression compared to Arc-AAV-*lacZ* infected cells (right panel of each pair, white arrows indicate ERα-negative, magenta arrows indicate ERα-positive infected cells). (B), representative reproductive cycles of mice that received Arc-AAV-*lacZ*, -*Esr1* g1 or g2; E, estrus, D, diestrus, P proestrus. Day 0 indicates the day of stereotaxic surgery. (C), pulsatile LH release in Arc-AAV-*lacZ*, -*Esr1* g1 or g2 mice, # indicate pulse detected by Cluster analysis(52). (D), mean±SEM days/week in proestrus before (from day −12 to day 14) and after infection (day 29 to day 56) in mice receiving Arc-AAV-*lacZ*, -*Esr1* g1 or g2. (E), individual values and mean±SEM LH pulses/h. (F), individual means and mean±SEM mean LH over the entire pretreatment sampling period. (G), mean±SEM LH before (con) and 15 min after kisspeptin (kiss) injection (left) and before (con) and 15 min after GnRH injection (right). * p<0.05

### Knockdown of ERα in arcuate kisspeptin neurons in adulthood increase ionotropic glutamatergic transmission to these cells but does not alter their short-term spontaneous firing rate

Arcuate kisspeptin neurons are postulated to form an interconnected network that is steroid sensitive and utilizes glutamatergic transmission at least in part for intranetwork communication (31). We thus hypothesized that loss of ERα specifically from arcuate kisspeptin neurons would increase their spontaneous firing rate and increase glutamatergic transmission to these cells, similar to what is observed in these cells in KERKO mice (20). As Arc-AAV-*Esr1* knockdown mice spend most time in estrus, similar to KERKO mice, we used estrus as the reproductive stage to examine the short-term (~10min) firing frequency of these neurons and AMPA-mediated excitatory glutamatergic postsynaptic currents (EPSCs). The firing frequency of Arc-AAV-*Esr1* infected cells was not different from Arc-AAV-*lacZ* infected cells (Fig 5A, B, Mann-Whitney U-test, p=0.14) even though there tends to be more cells firing at >1Hz in the Arc-AAV-*Esr1* group compared to the Arc-AAV-*lacZ* group (Fig 5C, Fisher’s exact test, *Esr1* vs *lacZ*, p=0.07). In contrast, the frequency and amplitude of glutamatergic EPSCs in arcuate kisspeptin cells in Arc-AAV-*Esr1* infected mice was greater than in the Arc-AAV-*lacZ* group (Fig 5D, E, F, two-tailed unpaired Student’s t-test, frequency p=0.0007, amplitude p=0.014).

**Fig 5.**
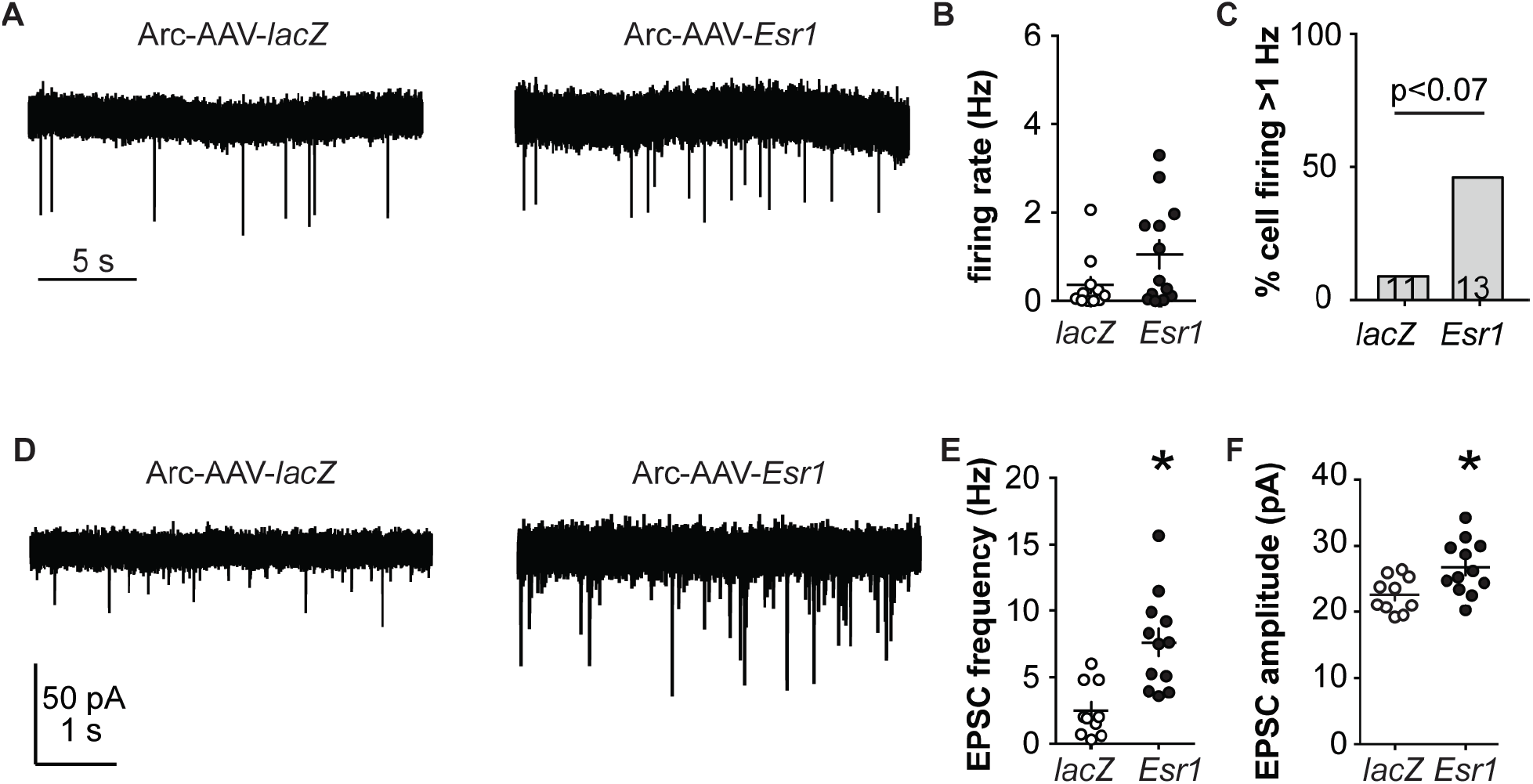
*Esr1* knockdown in arcuate kisspeptin neurons alters their cellular physiology. (A), representative extracellular recordings of firing rate. (B-C), individual values and mean±SEM firing rate (B) and percent of cells with firing rate >1 Hz (C); cells per group shown in bars. (D), representative whole-cell recordings of EPSCs. (E-F), individual values and mean±SEM of EPSC frequency (E) and amplitude (F). * p<0.05.

## Discussion

This study examined the roles of two hypothalamic kisspeptin neuronal populations in mediating estradiol feedback from cellular, molecular and whole-body physiology perspectives. We utilized both conventional kisspeptin-specific ERα knockout mice (KERKO) and CRISPR-Cas9 based viral vector-mediated knockdown of *Esr1*. The latter approach allows both temporal control and nucleus-specific manipulations to distinguish the role of ERα within each population in negative and positive feedback regulation of LH release and neurobiological properties.

AVPV kisspeptin neurons are postulated to convey estradiol positive feedback signals to generate the GnRH surge. Consistent with this postulate, these neurons are more excitable during positive feedback and also receive increased glutamatergic transmission (20, 25, 32, 33). AVPV kisspeptin cells in both KERKO and AVPV-AAV-*Esr1* models are less excitable compared to controls, firing fewer bursts and single action potentials in response to the same current injection. Results from the AVPV-AAV-*Esr1* model support and extend data from KERKO mice and provide evidence towards accepting the hypothesis that the role of ERα in shifting excitability is activational, independent of its role in the development of these cells (17).

Kisspeptin expression in AVPV cells is estradiol activated and fewer cells expressing *Kiss1* mRNA are detected in this region in KERKO mice (19). In AVPV-AAV-*Esr1* mice with adult knockdown, we also observed fewer cells express *Kiss1* mRNA compared to AVPV-AAV-*lacZ* mice, further supporting an activational role for estradiol in the adult physiology of these cells. The inability to sense estradiol through ERα in AVPV kisspeptin neurons may reduce production and release of kisspeptin and ultimately impair the downstream GnRH/LH surge. This could explain the blunted LH surges in AAV-*Esr1* injected mice. *Esr1* knockdown in AVPV, however, did not alter reproductive cyclicity monitored by changes in vaginal cytology, which reflect in part circulating steroids, in particular estradiol. In the AVPV-AAV-*Esr1* model, the blunted surges may be sufficient to trigger ovulation and to maintain cyclicity. Alternatively, cyclicity and the associated changes in sex steroids may be controlled by other cells that express ERα. Of interest, stress or neuronal androgen receptor KO disrupts the LH surge but without a change in estrous cyclicity (34, 35).

In support of the latter, several reproductive phenotypes of KERKO and AVPV-targeted ERα knockdown mice are different. KERKO mice tend to exhibit prolonged vaginal cornification and enlarged uteri, neither of which were observed in mice in which AAV-*Esr1* infection was targeted to the AVPV (SI Appendix Fig S2). In contrast, prolonged estrus and enlarged uteri were observed in mice in which Arc-AAV-*Esr1* infection was targeted to arcuate kisspeptin neurons. These latter neurons have been postulated to play a role in generating episodic GnRH output. Changes in episodic GnRH frequency drive gonadotropins and thus follicle development and steroidogenesis, including the estradiol rise, which triggers positive feedback and changes in vaginal cytology. Long-term firing output of arcuate kisspeptin neurons in brain slices is episodic and steroid modulated (36), and activation of these cells *in vivo* generates a pulse of LH release (37). Further evidence comes from *Tac2*-specific ERα KO mice, in which ERα is primarily deleted from the arcuate, not the AVPV, kisspeptin population. These mice also exhibit prolonged vaginal cornification (19). We thus hypothesize that ERα in arcuate kisspeptin neurons contributes to maintaining pulsatile LH release and mediates central estradiol negative feedback.

Consistent with this postulate, partial (~65%) adult knockdown of ERα in these cells altered the reproductive cycle. As KERKO mice exhibit increased LH-pulse frequency, we were initially surprised we did not observe differences in pulse frequency or mean LH levels in mice receiving Arc-AAV-*Esr1*. This may be attributable to single housing conditions in the present experiment, which may make mice prone to stress despite more than four weeks of handling before sampling. It is also possible that the pulse frequency during diestrus differs between Arc-AAV-*Esr1* and Arc-AAV-*lacZ* mice. Despite this lack of statistical difference in LH-pulse frequency, ERα knockdown mice had markedly reduced response to IP injection of both kisspeptin and GnRH, similar to KERKO mice (20). This suggests loss of ERα function in arcuate kisspeptin neurons may disrupt GnRH neuronal response to kisspeptin and/or the pituitary response to LH. This could arise from a disruption of negative feedback leading to overstimulation and thus desensitization of the hypothalamo-pituitary-gonadal axis or blunting of the response to administered neuropeptides.

Dissection of the electrophysiological properties of arcuate kisspeptin neurons revealed that glutamatergic transmission to these neurons was elevated when ERα is knocked down. This indicates the connectivity of these cells remains plastic even after puberty. The observation that targeted reduction of ERα in arcuate kisspeptin neurons increases glutamatergic transmission further suggests estradiol-sensitive interconnections among these cells provide many of their glutamatergic inputs. Given this, it is intriguing that the short-term firing rate of these cells was not increased, although there was a strong trend towards a greater percent of higher-frequency cells. The lack of change in mean firing rate may reflect the partial deletion of ERα in this population, with lower firing rate being preserved in cells with ERα, and elevated EPSC frequency arising at least in part from the high firing cells. It is also possible that long-term firing patterns of these Arc-AAV-*Esr1* infected arcuate kisspeptin neurons, which may be associated with episodic neuroendocrine activity, are disrupted. These data support the idea that glutamatergic inputs to arcuate kisspeptin neurons play an important role on maintaining normal reproductive function.

Although the CRISPR-Cas9 based knockdown approach allows spatial and temporal control, it too has caveats. For example, sgRNAs may have off-target actions on other regions of the genome beyond the sites predicted by the design software (38). To address this, we independently tested two sgRNAs that target *Esr1* to address the possible off-target effects among groups. We did not observe any differences between *Esr1* guide1 and guide2 groups. This suggests the phenotypes observed are primarily attributable to the deletion of ERα. Because of the nature of the nonhomologous end joining repair machinery activated after CRISPR-Cas9-initiated cuts, *Esr1* gene editing in each cell varies. It is difficult to assess each individual neuron to test if mutations at other genes are potentially involved in changes biophysical properties. Despite these variables, in the present study the systemic and cellular phenotypes in *Esr1* guide1 vs guide2 infected mice were quite consistent.

In conclusion, utilizing CRISPR-Cas9 AAV, we were able to successfully knockdown ERα in specific populations of kisspeptin neurons in adult female mice. Knockdown in each population recapitulated part of the KERKO model and furthers our understanding the role ERα in that population in regulating estradiol feedback.

## Materials and Methods

### Chemicals were from Sigma Chemical Company unless noted

#### Animals

The University of Michigan Institutional Animal Care and Use Committee approved all procedures. Adult female mice (60-150 days) were used. Mice were provided with water and Harlan 2916 chow (VetOne) *ad libitum* and were held on a 14L:10D light cycle (lights on 0400 Eastern Standard Time). To delete ERα specifically from kisspeptin cells (20), mice with the *Cre* recombinase gene knocked-in after the *Kiss1* promoter (*Kiss1*-Cre mice) were crossed with mice with a floxed *Esr1* gene, which encodes ERα (ERα floxed mice)(19). The expression of Cre recombinase mediates deletion of ERα in kisspeptin cells (KERKO mice). To visualize kisspeptin neurons for recording, mice heterozygous for both Kiss-Cre and floxed ERα were crossed with Cre-inducible YFP mice. Crossing mice heterozygous for all three alleles yielded litters that contained some mice that were homozygous for floxed ERα and at least heterozygous for both *Kiss1*-Cre and YFP; these were used as KERKO mice. Littermates of KERKO mice with wild type *Esr1, Kiss1*-Cre YFP (heterozygous or homozygous for either Cre or YFP) were used as controls; no differences were observed among these controls and they were combined. To generate kisspeptin-specific S. pyogenes Cas9 (Cas9)-expressing mice, mice with the *Cre* recombinase gene knocked-in after the *Kiss1* promoter (*Kiss1*-Cre mice)(39) were crossed with Rosa26-floxed STOP-Cas9 knockin mice (Jackson Lab Stock# 024857).

KERKO mice have disrupted estrous cycles with persistently cornified vaginal cytology typical of estrus; we thus used females in estrus as controls. Estrous cycle stage was determined by vaginal lavage. To examine the role of circulating estradiol, mice were ovariectomized (OVX) under isoflurane anesthesia (Abbott) and were either simultaneously implanted with a Silastic (Dow-Corning) capsule containing 0.625 µg of estradiol suspended in sesame oil (OVX+E) or not treated further (OVX)(24). Bupivacaine (0.25%, APP Pharmaceuticals) was provided local to incisions as an analgesic. These mice were studied 2-3d after surgery. Mice for electrophysiology were sacrificed at the time of estradiol positive feedback in the late afternoon (24). For free-floating immunochemistry staining, mice were perfused at 1700 EST 2-3d post OVX+E surgery at the expected peak of the estradiol-induced LH surge.

#### sgRNA design

For Cas9 target selection and generating single guide RNAs (sgRNA), 20-nt target sequences were selected to precede a 5’NGG protospacer-adjacent motif (PAM) sequence. To minimize off-targeting effects and maximize sgRNA activity, two CRISPR design tools were used to evaluate sgRNAs (27, 40) targeting mouse *Esr1* exon1. The two best candidates were selected based on lowest predicted off-target effects and highest activity. The target sequence for guide 1 is 5’-CACTGTGTTCAACTACCCCG-3’ (referred to as g1) and the target sequence for guide 2 is 3’-CTCGGGGTAGTTGAACACAG-5’ (referred to as g2). Because g1 and g2 were similarly effective in *Esr1* knockdown and effects on cycles, mice were combined for physiology studies. Control sgRNA sequence was designed to target *lacZ* gene from *Escherichia coli* (target sequence: 5’-TGCGCAGCCTGAATGGCGAA-3’).

#### In vitro validation of sgRNAs

C2C12 mouse myoblast cells (generous gift of Dr. Daniel Michele, University of Michigan) were grown in DMEM containing 10% FBS (Thermo Fisher) at 37°C in 5% CO2. Each individual sgRNA was introduced to BsmBI site of the lentiCRISPRv2 construct. Cells were co-transfected with one of the lentiCRISPRv2 plasmids containing sgRNAs and a standard GFP plasmid construct (41) using Lipofectamine 3000 (Invitrogen) according to the manufacturer’s instructions. Cells were selected for ~4 weeks with medium containing 1 μg/mL puromycin. Selected cells were harvested, DNA isolated using the Qiagen DNA Extraction Kit, and sequenced with primers for *Esr1*.

#### AAV vector production

To construct the AAV plasmid, a mCherry-U6 promoter-sgRNA scaffold segment was synthesized by Integrated DNA Technologies (IDT). After PCR amplification, the ligation product containing mCherry-U6 promoter-sgRNA scaffold was cloned in reverse orientation into a *Cre*-inducible AAV vector backbone (42). The individual sgRNAs (with an extra G added to the 5’-end of each sgRNA to increase guide efficiency (40)) were then inserted into a designed SapI site between U6 promoter and sgRNA scaffold component. AAV8 viral stocks were prepared at University of North Carolina Vector Core.

#### Stereotaxic injections

*Kiss1*Cre/Cas9-GFP female animals (>2 mo) were checked for estrous cycles for >10 days before surgery; only mice with regular 4-5 day cycles were used. Mice were anesthetized with 1.5%–2% isoflurane. AVPV injections were targeted to 0.55 mm posterior to Bregma, ±0.2mm lateral to midline, and 4.7 and 4.8mm ventral to dura. Arcuate injections were targeted to 1.5-1.7mm posterior to Bregma, ±0.2mm lateral to midline, and 5.9mm ventral to dura. 100nl virus injected bilaterally at the target coordinates at ~5nl/min. Estrous cycle monitoring continued after surgery for up to eight weeks.

#### Perfusion and free-floating immunohistochemistry

Mice were anesthetized with isoflurane and then transcardially perfused with PBS (15-20mL) then 10% neutral-buffered formalin for 10min (~50mL). Brains were placed into the same fixative overnight, followed by 30% sucrose for ≥24h for cryoprotection. Sections (30μm, 4 series) were cut on a cryostat (Leica CM3050S) and stored at −20°C in antifreeze solution (25% ethylene glycol, 25% glycerol in PBS). Sections were washed with PBS, treated with 0.1% hydrogen peroxide, and then placed in blocking solution (PBS containing 0.1% TritonX-100, 4% normal goat serum) for 1h at room temperature, then incubated with rabbit anti-ERα (#06-935, Millipore, 1:10,000) in blocking solution 48h at 4°C. Sections were washed then incubated with biotinylated anti-rabbit antibody (Jackson Immunoresearch, 1:500) followed by ABC amplification (Vector Laboratories, 1:500) and nickel-enhanced diaminobenzidine (Thermo Scientific) reaction (4.5min). Sections were washed with PBS and incubated overnight with chicken anti-GFP (ab13970, Abcam, 1:2000) and rat anti-mCherry (M11217, Invitrogen, 1:5000) in blocking solution. The next day, sections were washed and incubated with Alexa 594-conjugated anti-rat and Alexa 488-conjugated anti-chicken antibodies for 1h at room temperature (Molecular Probes, 1:500). Sections were mounted and coverslipped (VWR International 48393 251). Images were collected on a Zeiss AXIO Imager M2 microscope, and the number of immunoreactive GFP only, GFP/mCherry, and GFP/mCherry/ERα cells were counted in the injected region. The other kisspeptin region in the hypothalamus was examined and no infection of kisspeptin cell bodies was observed.

#### Brain slice preparation

All solutions were bubbled with 95%O_2_ and 5%CO_2_ for ≥15min before exposure to tissue and throughout experiments. Brains were rapidly removed 1.5-2h before lights off and placed in ice-cold sucrose saline solution containing (in mM): 250 sucrose, 3.5 KCl, 25 NaHCO_3_, 10 D-glucose, 1.25 Na_2_HPO_4_, 1.2 MgSO_4_, and 3.8 MgCl_2_. Coronal slides (300µm) were made with a Leica VT1200S. Slices were incubated in a 1:1 mixture of sucrose-saline and artificial cerebrospinal fluid (ACSF) containing (in mM): 135 NaCl, 3.5 KCl, 26 NaHCO_3_, 10 D-glucose, 1.25 Na_2_HPO_4_, 1.2 MgSO_4_, 2.5 CaCl_2_ for 30 min at room temperature. Slices were then transferred to 100% ACSF at room temperature for ≥30min before recording. Slices were used within 6h of preparation.

#### Electrophysiology recordings

Slices were transferred to a recording chamber and perfused with oxygenated ACSF (3mL/min) and heated by an in-line heater (Warner Instruments) to 30±1 °C. GnRH-GFP neurons were identified by brief illumination at 470nm using an upright fluorescence microscope Olympus BX51W1. Recording pipettes were pulled from borosilicate glass (type 7052, 1.65mm outer diameter and 1.12mm inner diameter; World Precision Instruments, Inc.) using a P-97 puller (Sutter Instruments) to obtain pipettes with a resistance of 2-3.5MW. Recordings were performed with an EPC-10 dual-patch clamp amplifier and Patchmaster acquisition software (HEKA Elektronik). Recorded cells were mapped to a brain atlas(43) to determine if cell location was related to response to treatment. No such correlation was observed in this study.

#### Extracellular recordings

Extracellular recordings were used to characterize firing rate as they maintain internal milieu and have minimal impact neuronal firing rate(44, 45). Recordings were made with receptors for ionotropic GABA_A_ and glutamate synaptic transmission antagonized (100µM picrotoxin, 20µM APV [D-(−)-2-amino-5-phosphonovaleric acid], 10µM CNQX [6-cyano-7-nitroquinoxaline]). Pipettes were filled with HEPES-buffered solution containing (in mM): 150 NaCl, 10 HEPES, 10 D-glucose, 2.5 CaCl_2_, 1.3 MgCl2, and 3.5 KCl (pH=7.4, 310 mOsm), and low-resistance (22±3MΩ) seals formed between the pipette and neuron after first exposing the pipette to the slice tissue in the absence of positive pressure. Recordings were made in voltage-clamp mode (0mV pipette holding potential) and signals acquired at 20kHz and filtered at 10kHz. Resistance of the loose seal was checked frequently during the first 3min of recordings to ensure a stable baseline, and also before and after a subsequent 10-min recording period; data were not use if seal resistance changed >30% or was >25MΩ. The first 5min of this 10-min recording were consistently stable among cells and were thus used for analysis.

#### Whole-cell recordings

For whole-cell patch-clamp recordings, three different pipette solutions were used depending on the goal. Most recordings were done with a physiologic pipette solution containing (in mM): 135 K gluconate, 10 KCl, 10 HEPES, 5 EGTA, 0.1 CaCl_2_, 4 MgATP and 0.4 NaGTP, pH 7.2 with NaOH, 302±3 mOsm. A similar solution containing 10mM neurobiotin was adjusted to similar osmolarity. A solution in which cesium gluconate replaced potassium gluconate was used to reduce potassium currents and allow better isolation of calcium currents. Membrane potentials reported were corrected online for liquid junction potential of −15.7 mV, same among all solutions(46).

After achieving a minimum 1.6GΩ seal and the whole-cell configuration, membrane potential was held at −70mV between protocols during voltage-clamp recordings. Series resistance (R_s_), input resistance (R_in_), holding current (I_hold_) and membrane capacitance (C_m_) were frequently measured using a 5mV hyperpolarizing step from −70mV (mean of 16 repeats). Only recordings with R_in_ >500 MΩ, I_hold_ −40 to 10pA and R_S_ <20MΩ, and stable C_m_ were accepted. R_s_ was further evaluated for stability and any voltage-clamp recordings with ∆R_s_ >15% were excluded; current-clamp recordings with ∆R_s_ >20% were excluded. There was no difference in I_hold_ or R_s_ among any comparisons.

For current-clamp recordings, depolarizing and hyperpolarizing current injections (−50 to +50pA, 500ms, 10pA increments) were applied from an initial membrane potential of −71±2 mV, near the resting membrane potential of these cells (47).

For voltage-clamp recordings of excitatory postsynaptic currents (EPSCs), membrane potential was held at −68mV, the reversal potential for GABA_A_-receptor mediated currents, and ACSF contained picrotoxin (100μM), and APV (D-(−)-2-amino-5-phosphonovaleric acid, 20μM).

For voltage-clamp recordings of I_T_, ACSF containing antagonists of ionotropic GABA_A_ and glutamate receptors was supplemented with TTX (2µM) and the Cs-based pipette solution was used. Two voltage protocols were used to isolate I_T_ as reported (25). First, total calcium current activation was examined. Inactivation was removed by hyperpolarizing the membrane potential to −110mV for 350ms (not shown in figures). Next a 250ms prepulse of −110mV was given. Then membrane potential was varied in 10mV increments for 250ms from −110 to −30mV. Finally, test pulse of −40mV for 250ms was given. From examination of the current during the test pulse, it was evident that no sustained (high-voltage activated) calcium current was activated at potentials more hyperpolarized than −40mV. To remove HVA contamination from the step to −30mV, a second protocol was used in which removal of inactivation (−110mV, 350ms) was followed by a 250ms prepulse at −40mV, then a step for 250ms at −30mV and finally a test pulse of −40mV for 250ms. I_T_ was isolated by subtracting the trace following the −40mV prepulse from those obtained after the −110mV prepulse for the depolarized variable step to −30mV; raw traces from the initial voltage protocol were used without subtraction for variable steps from −110mV to −40mV because of the lack of observed activation of HVA at these potentials. Activation of I_T_ was assessed from the resulting family of traces by peak current during the variable step phase. Inactivation of I_T_ was assessed from the peak current during the final −40mV test pulse.

#### Post hoc identification of ERα

The pipette solution containing neurobiotin was used for recordings cells from AAV-injected mice. An outside-out patch was formed after recording to reseal the membrane and the location of cells was marked on a brain atlas (43). The brain slices were fixed overnight in 10% formalin at 4°C and changed to PBS. Slices were photo-bleached with a UV illuminator for ~72h and checked to ensure no visible fluorescent signal was observed. Slices were then placed in blocking solution for 1h, then incubated with rabbit anti-ERα for 48h at 4°C. Slices were washed and then incubated with Alexa 594-conjugated anti-rabbit and Alexa 350-conjugated neutravidin for 2h at room temperature (Molecular Probes, 1:500). Slices were mounted, coverslipped and imaged as above. Cells with neurobiotin-labeling were examined for ERα-immunoreactivity.

#### Single-cell PCR

Cells for single cell PCR were collected as previously described (48). Patch pipettes (2–3MΩ) were filled with 5–8μL of an RNase free solution containing (in mM) 135 K-gluconate, 10 KCl, 10 HEPES, 5 EGTA, 4.0 Mg-ATP, 0.4 Na-GTP, and 1.0 CaCl_2_ (pH 7.3, 305 mOsm). Additionally, before use 1U/µL Protector RNase Inhibitor (Roche) was added to the pipette solution. Single Cell RNA was harvested from the target cells in whole-cell configuration after current-clamp recordings; cytoplasm was aspirated into the pipette and expelled into a 0.2mL tube containing reverse transcriptase buffer (Superscript Vilo cDNA Synthesis Kit, Invitrogen/ThermoFisher), volume was adjusted to 20µL with molecular grade water. Cell contents were reverse transcribed following manufacturer’s instructions. False harvests, in which the pipette was lowered into the slice preparation but no aspiration of cell contents occurred, were used to estimate background contamination. These were performed on each recording day. Additionally, a standard curve of mouse hypothalamic RNA (1, 0.1, 0.01, 0.001 ng/μL final concentration) and a water blank (negative control) were reverse transcribed. Single-cell cDNA, controls, and the standard curve were preamplified for 15 cycles using TaqMan PreAmp Master Mix (Invitrogen/ThermoFisher) as previously described (49). Quantitative PCR was performed using 5μL of diluted preamplified DNA (1:10) per reaction, in duplicate, for 50 cycles (TaqMan Gene Expression Master Mix; Invitrogen). Single cell cDNA was assayed for: *Kiss1, Th, Esr1, Esr2, Pgr, Cacna1g, Cacna1h, Cacna1i, Hcn1 Hcn2 Hcn3 Hcn4; Syn1* was used as housekeeping gene; only Syn1 positive cells were analyzed. Single cells were considered positive for a transcript if their threshold was a minimum of three cycles earlier (8 fold greater) than the false harvests and the reverse transcribed and preamplified water blank sample. TaqMan PrimeTime qPCR assays for mRNAs (SI Appendix Table 2) were purchased from IDT.

#### *Tail-tip blood collection for LH* pulses

Ovary-intact *Kiss1*Cre-Cas9 adult female mice with AAV-*lacZ* and AAV-*Esr1* targeted to the arcuate nucleus were singly-housed were handled daily ≥4wks before sampling. Vaginal cytology of was determined for ≥10 days before sampling. As the majority of AAV-*Esr1* arcuate targeted mice (6 of 9) exhibit prolonged cornification typical of estrus, all mice (*Esr1* and *lacZ*) were sampled during estrus. Repetitive tail-tip blood collecting was performed as described (50). After the excision of the very tip of the tail, blood (6µL) was collected every 6 min for 2h from 1pm to 3pm. At the end of this frequent sampling period, mice received a single intraperitoneal injection of kisspeptin (65µg/kg)(51). Blood was collected just before and 15 min after kisspeptin injection. GnRH (150µg/kg)(49) was injected 40-45 min after kisspeptin, with blood collected immediately before and 15 min after GnRH injection.

#### Tail-tip blood collection for LH surge

Ovary-intact *Kiss1*Cre-Cas9 adult female mice with AAV-*lacZ* and AAV-*Esr1* targeted to the AVPV were singly-housed. Tail blood was collected as above on proestrus at 3, 4 and 5pm EST (lights are off at 5pm EST in the mouse room). One to two weeks later, these same mice were then subjected to OVX+E surgery and tail blood (6µL) was collected 2-3 days post-surgery at 9am and 5pm EST.

#### LH assay

Whole blood was immediately diluted in 54μL of 0.1M PBS with 0.05% Tween 20 and homogenized and kept on ice. Samples were stored at −20 °C for a subsequent ultrasensitive LH assay(50). Intraassay CV was 2.2%; interassay CVs were 7.3% (low QC, 0.13 ng/mL), 5.0% (medium QC, 0.8 ng/mL) and 6.5% (high QC, 2.3ng/mL). Functional sensitivity was 0.016ng/mL.

#### Data analysis and statistics

Data were analyzed offline using custom software written in IgorPro 6.31 (Wavemetrics). For targeted extracellular recordings, mean firing rate in Hz was determined over 5 min of stable recording. In experiments examining I_T_, the peak current amplitude at each step potential (V) was first converted to conductance using the calculated reversal potential of Ca^2+^ (E_Ca_) and G=I/(E_Ca_ - V), because driving force was linear over the range of voltages examined. The voltage dependencies of activation and steady-state inactivation were described with a single Boltzmann distribution: G(V)= G_max_/_-_(1- exp [(V_1/2_ - Vt)/k]), where G_max_ is the maximal conductance, V_1/2_ is the half-maximal voltage, and k is the slope. Current density of I_T_ at each tested membrane potential was determined by dividing peak current by membrane capacitance. LH pulses were detected by a version of Cluster (52) transferred to IgorPro using cluster sizes of two points for both peak and nadir and t-scores of two for detection of increases and decreases. Data were analyzed using Prism 7 (GraphPad Software) and reported as mean±SEM. The number of cells per group is indicated by n. For two by two designs, data were normally distributed and analyzed by two-way ANOVA or two-way repeated-measures (RM) with Holm-Sidak *post hoc* analysis. For two group comparisons normally distributed data were analyzed by two-tailed unpaired Student’s *t*-test; non-normal data were analyzed by Mann-Whitney U test. For categorical data, for more than 3 categories, Chi-square test of independence were used with Fisher’s exact test as *post hoc* analysis. For two categories, Fisher’s exact test was used. For each electrophysiological parameter comparison, no more than three cells per mouse was used in control and KERKO mice; no more than four cells per mouse was used for AAV-infected mice. No less than five mice were tested per parameter. The variance of the data was no smaller within an animal than among animals. For IHC staining, LH surge and LH pulse measurements, and reproductive cyclicity, at least three mice were tested per AAV vector.

## Acknowledgements

We thank Elizabeth Wagenmaker for expert technical assistance. We thank Dr. Daniel Michele for sharing C2C12 cell lines with us. We thank University of Virginia Ligand Core (NIH HD028934) for performing the ultra-sensitive LH assay. Supported by NIH HD41469 (SMM), DK095201 (YMS) and the Michigan Diabetes Research Center (MGM, P30DK020572).

## Supplementary Information for

This PDF file includes:

Supplementary Figs. S1 to S4

Table S1 to S2

**Fig. S1.**
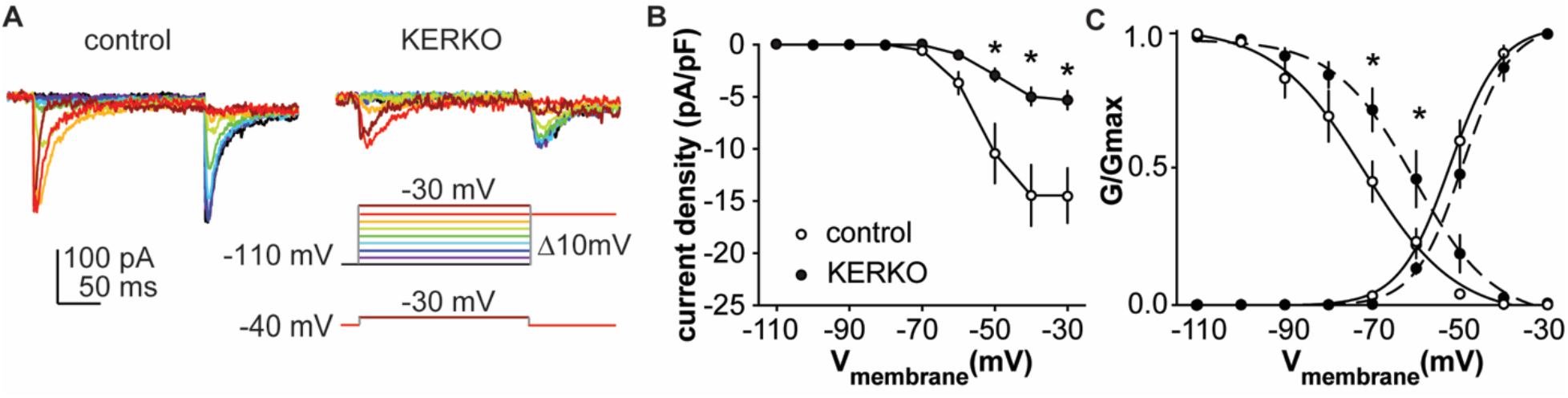
T-type calcium currents are reduced in AVPV kisspeptin neurons from KERKO compared to control mice. (A), voltage protocol (bottom right) and representative IT in control (left, n=8) and KERKO groups (right, n=7). (B), mean±SEM IT current density in control (white symbols, n=8) and KERKO groups (black symbols, n=7). (C), voltage dependence of IT conductance activation and inactivation in cells from control (n=8) and KERKO mice (n=7). * p<0.05.

**Fig S2.**
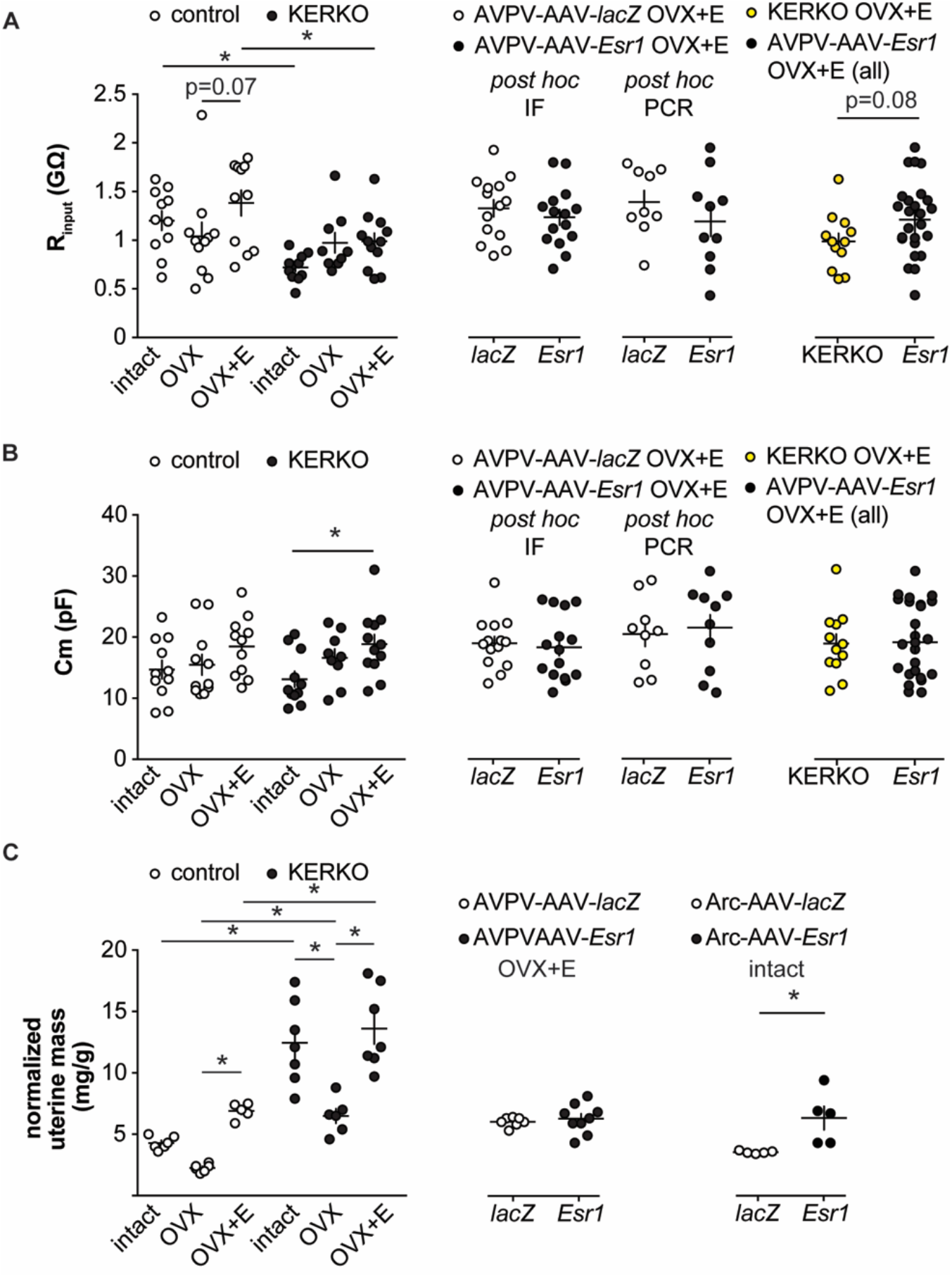
(A-B), individual values and mean±SEM input resistance (R_input_, A) and cell capacitance (Cm, B) for AVPV kisspeptin neurons in control and KERKO mice (left), in mice with AAV vector delivered to AVPV region (middle with *Esr1* status confirmed by either immunofluorescence (IF) single-cell qPCR (PCR) *post hoc)*, and in AVPV-AAV-*Esr1* infected mice (combined detection methods) vs KERKO mice (right). (C) individual values and mean±SEM normalized uterine mass (uterine mass/body mass, mg/g) of control and KERKO mice (left), of mice with AAV delivered to AVPV region, OVX+E (middle), and of mice with AAV delivered to arcuate region, intact (right). * p<0.05. The lack of a significant drop in the ratio is likely attributable to the short duration post OVX (2 days).

**Fig S3.**
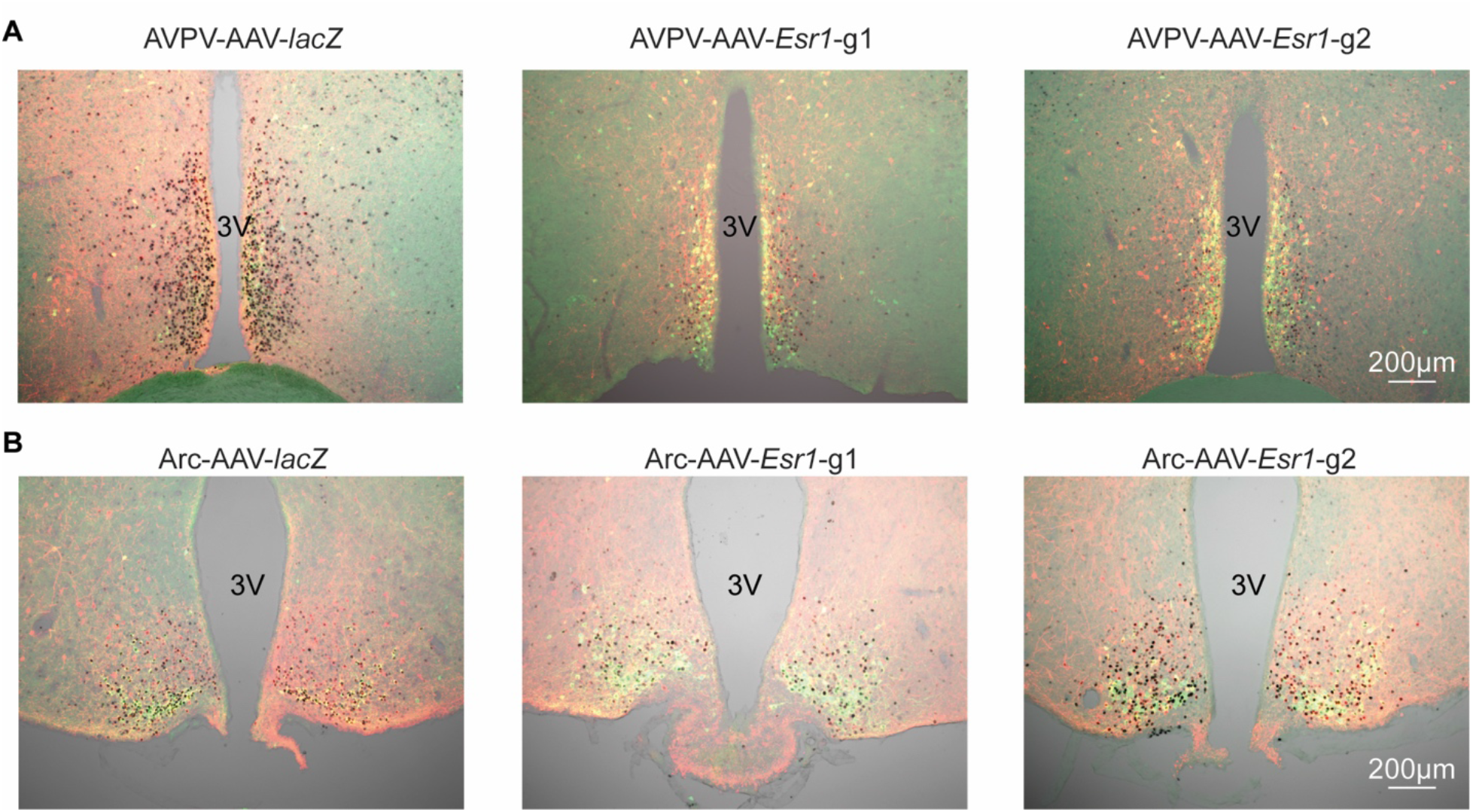
Bilateral delivery of AAV-lacZ, and AAV-Esr1 (g1 and g2) to AVPV (A) and arcuate (B) of adult female mice. Immunofluorescence was used to detect GFP (green), mCherry (red) and immunohistochemistry to detect ERα (black).

**Fig S4.**
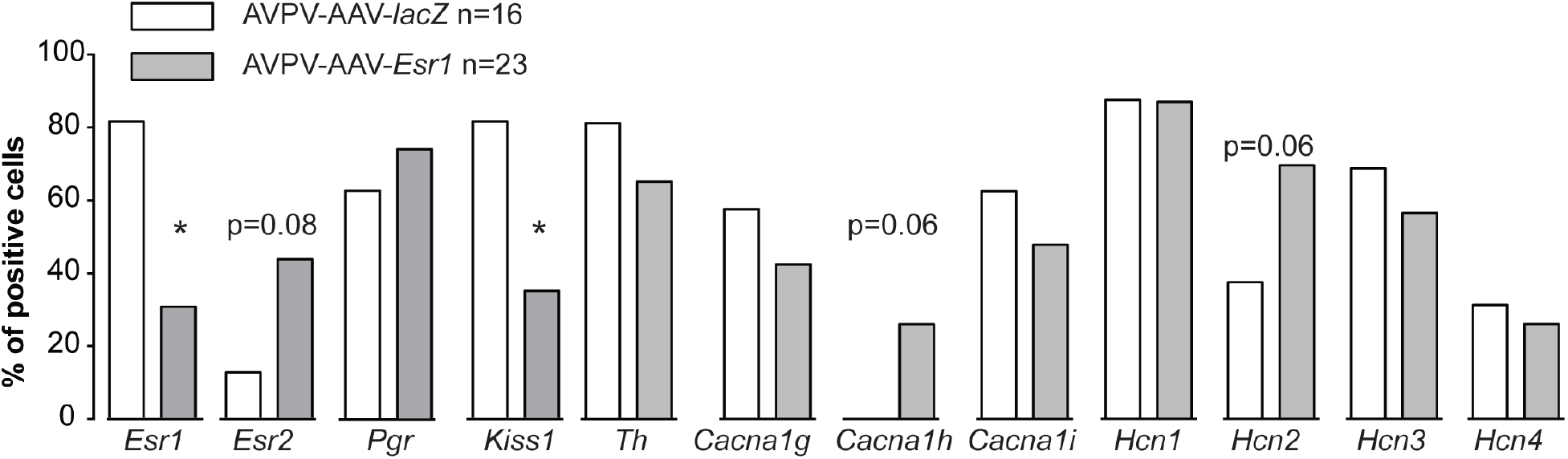
Single-cell qPCR for mRNA from AVPV kisspeptin neurons in mice with AAV vector delivered to AVPV region. Bar graphs show percentage of cell positive for each gene. * p<0.05, Fisher’s exact test.

**Table S1.**
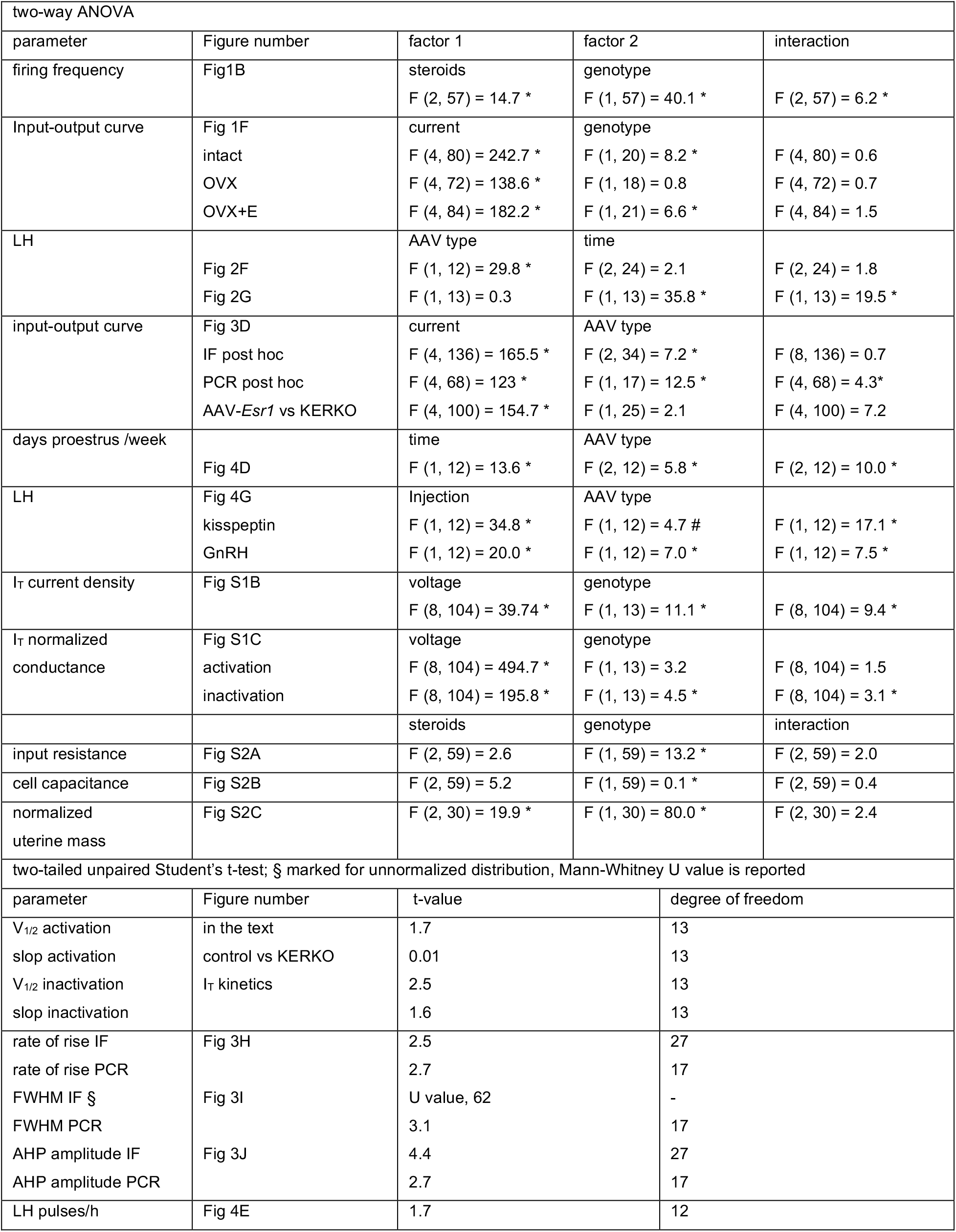
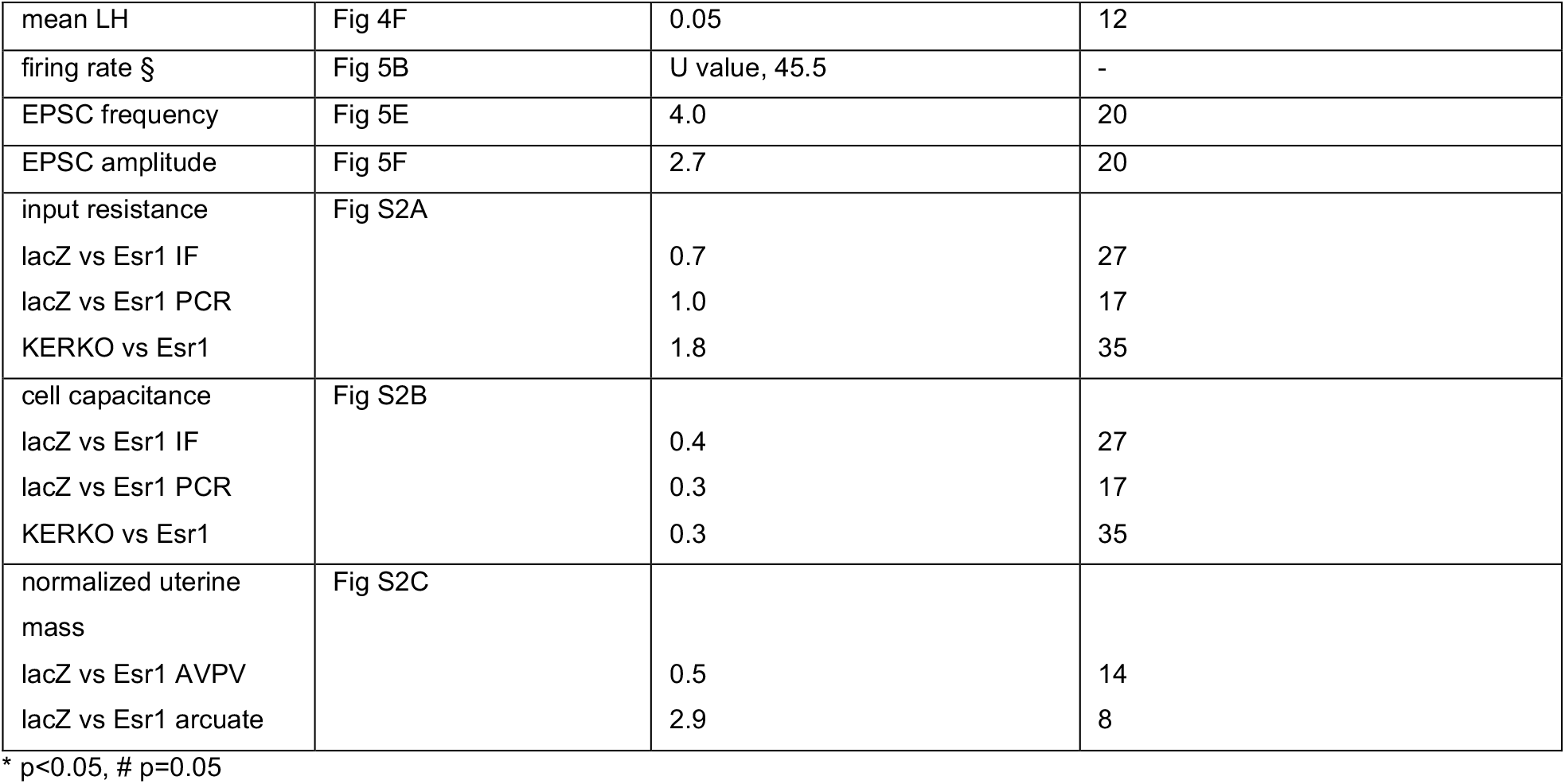
Statistical test parameters.

| two-way ANOVA |  |  |  |  |
| --- | --- | --- | --- | --- |
| parameter | Figure number | factor 1 | factor 2 | interaction |
| firing frequency | Fig1B | steroids<br>$F(2, 57) = 14.7^*$ | genotype<br>$F(1, 57) = 40.1^*$ | $F(2, 57) = 6.2^*$ |
| Input-output curve | Fig 1F<br>intact<br>OVX<br>OVX+E | current<br>$F(4, 80) = 242.7^*$<br>$F(4, 72) = 138.6^*$<br>$F(4, 84) = 182.2^*$ | genotype<br>$F(1, 20) = 8.2^*$<br>$F(1, 18) = 0.8$<br>$F(1, 21) = 6.6^*$ | $F(4, 80) = 0.6$<br>$F(4, 72) = 0.7$<br>$F(4, 84) = 1.5$ |
| LH | Fig 2F<br>Fig 2G | AAV type<br>$F(1, 12) = 29.8^*$<br>$F(1, 13) = 0.3$ | time<br>$F(2, 24) = 2.1$<br>$F(1, 13) = 35.8^*$ | $F(2, 24) = 1.8$<br>$F(1, 13) = 19.5^*$ |
| input-output curve | Fig 3D<br>IF post hoc<br>PCR post hoc<br>AAV-*Esr1* vs KERKO | current<br>$F(4, 136) = 165.5^*$<br>$F(4, 68) = 123^*$<br>$F(4, 100) = 154.7^*$ | AAV type<br>$F(2, 34) = 7.2^*$<br>$F(1, 17) = 12.5^*$<br>$F(1, 25) = 2.1$ | $F(8, 136) = 0.7$<br>$F(4, 68) = 4.3^*$<br>$F(4, 100) = 7.2$ |
| days proestrus /week | Fig 4D | time<br>$F(1, 12) = 13.6^*$ | AAV type<br>$F(2, 12) = 5.8^*$ | $F(2, 12) = 10.0^*$ |
| LH | Fig 4G<br>kisspeptin<br>GnRH | Injection<br>$F(1, 12) = 34.8^*$<br>$F(1, 12) = 20.0^*$ | AAV type<br>$F(1, 12) = 4.7^{\#}$<br>$F(1, 12) = 7.0^*$ | $F(1, 12) = 17.1^*$<br>$F(1, 12) = 7.5^*$ |
| $I_T$ current density | Fig S1B | voltage<br>$F(8, 104) = 39.74^*$ | genotype<br>$F(1, 13) = 11.1^*$ | $F(8, 104) = 9.4^*$ |
| $I_T$ normalized conductance | Fig S1C<br>activation<br>inactivation | voltage<br>$F(8, 104) = 494.7^*$<br>$F(8, 104) = 195.8^*$ | genotype<br>$F(1, 13) = 3.2$<br>$F(1, 13) = 4.5^*$ | $F(8, 104) = 1.5$<br>$F(8, 104) = 3.1^*$ |
|  |  | steroids | genotype | interaction |
| input resistance | Fig S2A | $F(2, 59) = 2.6$ | $F(1, 59) = 13.2^*$ | $F(2, 59) = 2.0$ |
| cell capacitance | Fig S2B | $F(2, 59) = 5.2$ | $F(1, 59) = 0.1^*$ | $F(2, 59) = 0.4$ |
| normalized uterine mass | Fig S2C | $F(2, 30) = 19.9^*$ | $F(1, 30) = 80.0^*$ | $F(2, 30) = 2.4$ |
| two-tailed unpaired Student's t-test; § marked for unnormalized distribution, Mann-Whitney U value is reported |  |  |  |  |
| parameter | Figure number | t-value | degree of freedom |  |
| $V_{1/2}$ activation | in the text | 1.7 | 13 | |
| slop activation | control vs KERKO | 0.01 | 13 |  |
| $V_{1/2}$ inactivation | $I_T$ kinetics | 2.5 | 13 | |
| slop inactivation |  | 1.6 | 13 |  |
| rate of rise IF | Fig 3H | 2.5 | 27 |  |
| rate of rise PCR |  | 2.7 | 17 |  |
| FWHM IF § | Fig 3I | U value, 62 | - |  |
| FWHM PCR |  | 3.1 | 17 |  |
| AHP amplitude IF | Fig 3J | 4.4 | 27 |  |
| AHP amplitude PCR |  | 2.7 | 17 |  |
| LH pulses/h | Fig 4E | 1.7 | 12 |  |

|  |  |  |  |
| --- | --- | --- | --- |
| mean LH | Fig 4F | 0.05 | 12 |
| firing rate § | Fig 5B | U value, 45.5 | - |
| EPSC frequency | Fig 5E | 4.0 | 20 |
| EPSC amplitude | Fig 5F | 2.7 | 20 |
| input resistance | Fig S2A |  |  |
| lacZ vs Esr1 IF |  | 0.7 | 27 |
| lacZ vs Esr1 PCR |  | 1.0 | 17 |
| KERKO vs Esr1 |  | 1.8 | 35 |
| cell capacitance | Fig S2B |  |  |
| lacZ vs Esr1 IF |  | 0.4 | 27 |
| lacZ vs Esr1 PCR |  | 0.3 | 17 |
| KERKO vs Esr1 |  | 0.3 | 35 |
| normalized uterine mass | Fig S2C |  |  |
| lacZ vs Esr1 AVPV |  | 0.5 | 14 |
| lacZ vs Esr1 arcuate |  | 2.9 | 8 |

\* p<0.05, # p=0.05

**Table S2.**
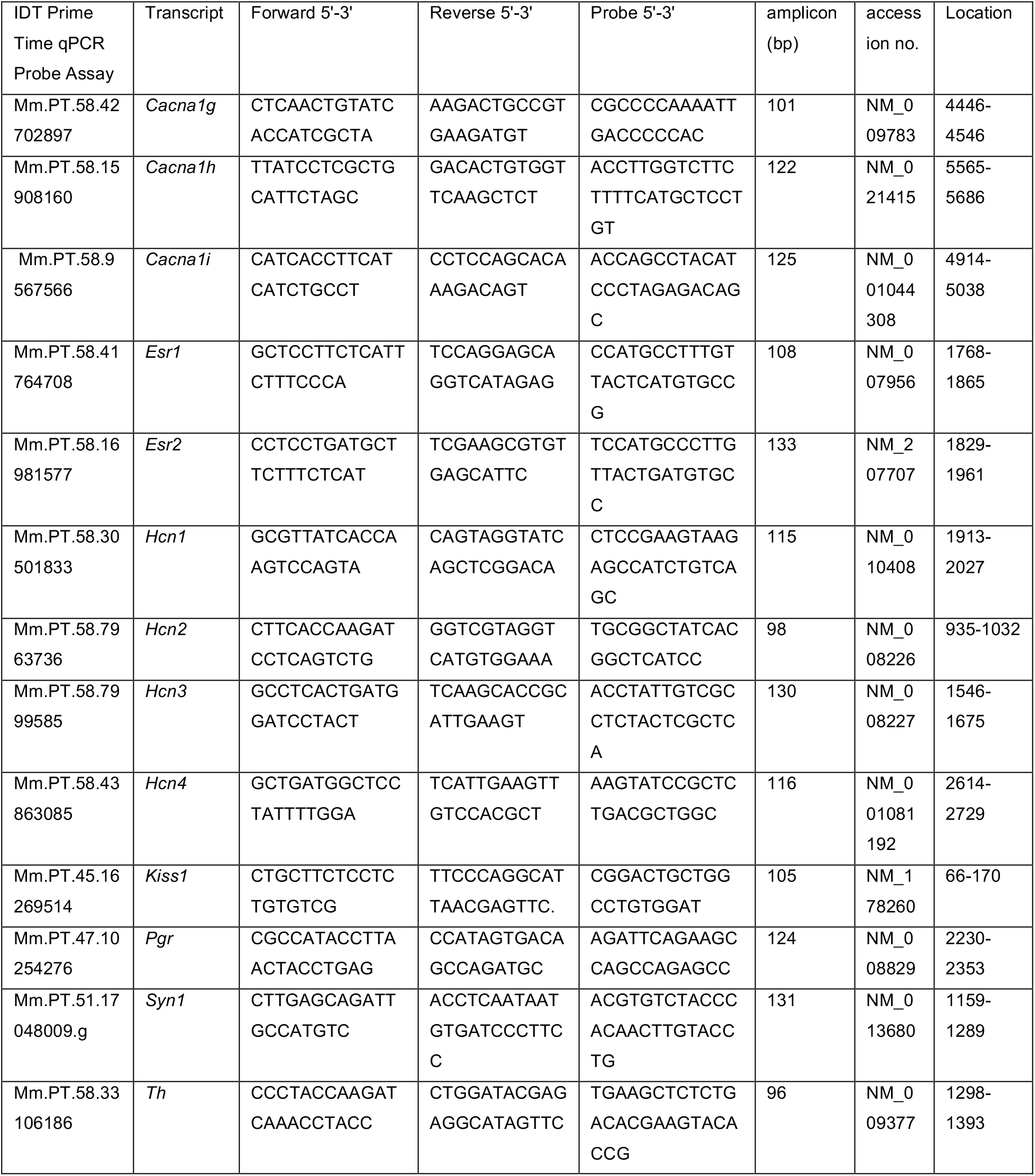
Primer probes used for single-cell qPCR.

## References

1. Macaluso, M, et al. (2010) A public health focus on infertility prevention, detection, and management. Fertil Steril 93(1):16.e1-10.

2. Plant, TM, Zeleznik AJ (2015) Knobil and Neill’s physiology of reproduction 4th edition (Elsevier). 4thEdition Ed.

3. Helm, KD, Nass, RM, Evans, WS, Nass, RM, Evans WS (2009) Physiologic and Pathophysiologic Alterations of the Neuroendocrine Components of the Reproductive Axis. Yen & Jaffe’s Reproductive Endocrinology (Elsevier), pp 441–488.

4. Christian, CA, Glidewell-Kenney, C, Jameson, JL, Moenter SM (2008) Classical estrogen receptor α signaling mediates negative and positive feedback on gonadotropin-releasing hormone neuron firing. Endocrinology 149(May):5328–5334.

5. Glanowska, KM, Venton, BJ, Moenter SM (2012) Fast scan cyclic voltammetry as a novel method for detection of real-time gonadotropin-releasing hormone release in mouse brain slices. J Neurosci 32(42):14664–9.

6. Moenter, SM, Caraty, A, Karsch FJ (1990) The estradiol-induced surge of gonadotropin-releasing hormone in the ewe. Endocrinology 127(3):1375–84.

7. Christian, CA, Moenter SM (2010) The neurobiology of preovulatory and estradiol-induced gonadotropin-releasing hormone surges. Endocr Rev 31(4):544–77.

8. Hrabovszky, E, et al. (2001) Estrogen receptor-beta immunoreactivity in luteinizing hormone-releasing hormone neurons of the rat brain. Endocrinology 142(7):3261–4.

9. Oakley, AE, Clifton, DK, Steiner RA (2009) Kisspeptin signaling in the brain. Endocr Rev 30(October):713–743.

10. Lehman, MN, Coolen, LM, Goodman RL (2010) Minireview: Kisspeptin/neurokinin B/dynorphin (KNDy) cells of the arcuate nucleus: a central node in the control of gonadotropin-releasing hormone secretion. Endocrinology 151(8):3479–3489.

11. Kumar, D, et al. (2015) Specialized subpopulations of kisspeptin neurons communicate with GnRH neurons in female mice. Endocrinology 156(1):32–38.

12. Yip, SH, Boehm, U, Herbison, AE, Campbell RE (2015) Conditional viral tract tracing delineates the projections of the distinct kisspeptin neuron populations to gonadotropin-releasing hormone (GnRH) neurons in the mouse. Endocrinology 156(7):2582–94.

13. Smith, JT, Cunningham, MJ, Rissman, EF, Clifton, DK, Steiner RA (2005) Regulation of Kiss1 gene expression in the brain of the female mouse. Endocrinology 146(9):3686–3692.

14. Pielecka-Fortuna, J, Chu, Z, Moenter SM (2008) Kisspeptin acts directly and indirectly to increase gonadotropin-releasing hormone neuron activity and its effects are modulated by estradiol. Endocrinology 149(4):1979–86.

15. Messager, S, et al. (2005) Kisspeptin directly stimulates gonadotropin-releasing hormone release via G protein-coupled receptor 54. Proc Natl Acad Sci U S A 102(5):1761–1766.

16. Han, S, et al. (2005) Activation of gonadotropin-releasing hormone neurons by kisspeptin as a neuroendocrine switch for the onset of puberty. J Neurosci 25(49):11349–56.

17. Mayer, C, et al. (2010) Timing and completion of puberty in female mice depend on estrogen receptor alpha-signaling in kisspeptin neurons. Proc Natl Acad Sci U S A 107(52):22693–22698.

18. Dubois, SL, et al. (2015) Positive, but not negative feedback actions of estradiol in adult female mice require estrogen receptor α in kisspeptin neurons. Endocrinology (Mar):156(3):1111-20.

19. Greenwald-Yarnell, ML, et al. (2016) ERa in Tac2 neurons regulates puberty onset in female mice. Endocrinology 157(4):1555–1565.

20. Wang, L, Burger, LL, Greenwald-Yarnell, ML, Myers, MG, Moenter SM (2018) Glutamatergic transmission to hypothalamic kisspeptin neurons is differentially regulated by estradiol through estrogen receptor α in adult female mice. J Neurosci 38(5):1061 LP-1072.

21. Semaan, SJ, et al. (2010) BAX-dependent and BAX-independent regulation of Kiss1 neuron development in mice. Endocrinology 151(12):5807–5817.

22. Kumar, D, et al. (2014) Murine arcuate nucleus kisspeptin neurons communicate with GnRH neurons in utero. J Neurosci 34(10).

23. Swiech, L, et al. (2014) In vivo interrogation of gene function in the mammalian brain using CRISPR-Cas9. Nat Biotechnol 33(1):102–106.

24. Christian, CA, Mobley, JL, Moenter SM (2005) Diurnal and estradiol-dependent changes in gonadotropin-releasing hormone neuron firing activity. Proc Natl Acad Sci U S A 102(43):15682–7.

25. Wang, L, DeFazio, RA, Moenter SM (2016) Excitability and burst generation of AVPV kisspeptin neurons are regulated by the estrous cycle via multiple conductances modulated by estradiol action. eNeuro 3(3):e0094-16.

26. Lee, JH, Gomora, JC, Cribbs, LL, Perez-Reyes E (1999) Nickel block of three cloned T-type calcium channels: low concentrations selectively block alpha1H. Biophys J 77(6):3034–42.

27. Ran, FA, et al. (2013) Genome engineering using the CRISPR-Cas9 system. Nat Protoc 8(11):2281–2308.

28. Milanesi, L, de Boland, AR, Boland R (2008) Expression and localization of estrogen receptor α in the C2C12 murine skeletal muscle cell line. J Cell Biochem 104(4):1254–1273.

29. Sanjana, NE, Shalem, O, Zhang F (2014) Improved vectors and genome-wide libraries for CRISPR screening. Nat Methods 11(8):783–784.

30. Hilton, HN, Graham, JD, Clarke CL (2015) Minireview: progesterone regulation of proliferation in the normal human breast and in breast cancer: a tale of two scenarios? Mol Endocrinol 29(9):1230–1242.

31. Qiu, J, et al. (2016) High-frequency stimulation-induced peptide release synchronizes arcuate kisspeptin neurons and excites GnRH neurons. Elife 5(AUGUST):e16246.

32. Piet, R, et al. (2013) Estrous cycle plasticity in the hyperpolarization-activated current Ih Is mediated by circulating 17-estradiol in preoptic area kisspeptin neurons. J Neurosci 33(26):10828–10839.

33. Zhang, C, et al. (2013) Molecular mechanisms that drive estradiol-dependent burst firing of Kiss1 neurons in the rostral periventricular preoptic area. Am J Physiol Endocrinol Metab 305(11):E1384-97.

34. Wagenmaker, ER, Moenter SM (2017) Exposure to acute psychosocial stress disrupts the luteinizing hormone surge independent of estrous cycle alterations in female mice. Endocrinology 158(8):2593–2602.

35. Walters, KA, et al. (2018) The Role of Central Androgen Receptor Actions in Regulating the Hypothalamic-Pituitary-Ovarian Axis. Neuroendocrinology 106(4):389–400.

36. Vanacker, C, Moya, MR, DeFazio, RA, Johnson, ML, Moenter SM (2017) Long-term recordings of arcuate nucleus kisspeptin neurons reveal patterned activity that is modulated by gonadal steroids in male mice. Endocrinology. doi:10.1210/en.2017-00382.

37. Clarkson, J, et al. (2017) Definition of the hypothalamic GnRH pulse generator in mice. Proc Natl Acad Sci 114(47):E10216–E10223.

38. Anderson, KR, et al. (2018) CRISPR off-target analysis in genetically engineered rats and mice. Nat Methods 15(7):512–514.

39. Cravo, RM, et al. (2011) Characterization of Kiss1 neurons using transgenic mouse models. Neuroscience 173:37–56.

40. Doench, JG, et al. (2014) Rational design of highly active sgRNAs for CRISPR-Cas9–mediated gene inactivation. Nat Biotechnol 32(12):1262–1267.

41. Ramakrishnan, SK, et al. (2016) HIF2α Is an Essential Molecular Brake for Postprandial Hepatic Glucagon Response Independent of Insulin Signaling. Cell Metab 23(3):505–516.

42. Flak, JN, et al. (2017) A leptin-regulated circuit controls glucose mobilization during noxious stimuli. J Clin Invest 127(8):3103–3113.

43. Paxinos, G, Franklin K (2001) The Mouse Brain in Stereotaxic Coordinates: Second Edition. (Elsevier Academic Press, San Diego, CA).

44. Nunemaker, CS, DeFazio, RA, Moenter SM (2003) A targeted extracellular approach for recording long-term firing patterns of excitable cells: a practical guide. Biol Proced Online 5:53–62.

45. Alcami, P, Franconville, R, Llano, I, Marty A (2012) Measuring the firing rate of high-resistance neurons with cell-attached recording. J Neurosci 32(9):3118–30.

46. Barry PH (1994) JPCalc, a software package for calculating liquid junction potential corrections in patch-clamp, intracellular, epithelial and bilayer measurements and for correcting junction potential measurements. J Neurosci Methods 51(1):107–16.

47. DeFazio, RA, Elias, CF, Moenter SM (2014) GABAergic transmission to kisspeptin neurons is differentially regulated by time of day and estradiol in female mice. J Neurosci 34(49):16296–308.

48. Ruka, KA, Burger, LL, Moenter SM (2013) Regulation of arcuate neurons coexpressing kisspeptin, neurokinin, B, and dynorphin by modulators of neurokinin 3 and κ-opioid receptors in adult male mice. Endocrinology 154(8):2761–71.

49. Glanowska, KM, Burger, LL, Moenter SM (2014) Development of Gonadotropin-releasing hormone secretion and pituitary response. J Neurosci 34(45).

50. Steyn, FJ, et al. (2013) Development of a methodology for and assessment of pulsatile luteinizing hormone secretion in juvenile and adult male mice. Endocrinology 154(12):4939–4945.

51. Hanchate, NK, et al. (2012) Kisspeptin-GPR54 signaling in mouse NO-synthesizing neurons participates in the hypothalamic control of ovulation. J Neurosci 32(3).

52. Veldhuis, JD, Johnson ML (1986) Cluster analysis: a simple, versatile, and robust algorithm for endocrine pulse detection. Am J Physiol 250(4 Pt 1):E486-93.

